# How specific structural differences of Bcl2 proteins modulate the interaction with BH3 domains and apoptotic function

**DOI:** 10.64898/2026.07.17.739112

**Authors:** Anton Hanke, Christina Elsner, Simone Aureli, Enrica Bordignon, Francesco Luigi Gervasio

**Author notes:** AH conducted experiments and interpreted data. AH, EB, FLG devised study design. AH wrote and AH, CE, SA, EB, FLG revised and finalized the manuscript. FLG and EB acquired funding.

## Abstract

Intrinsic apoptosis is mainly regulated through a network of conserved interactions between Bcl-2 proteins involving hydrophobic binding grooves and BH3 domains. Despite these conserved interfaces, family members exhibit distinct binding affinities and play opposing roles in apoptosis. While static structural differences partially account for this divergence, it remains unclear how opposing apoptotic function reflects in BH3 helix engagement of individual members. Here, we investigate how a BidBH3 peptide engages with the hydrophobic groove of full-length membrane-anchored Bcl-xL and Bax to identify shared and unique features of binding that may relate to distinct apoptotic functions. Using state-of-the-art enhanced-sampling simulations, we mapped the complete binding and folding landscapes of these critical cell-death regulators in membranes. Our simulations align with experimental measurements in terms of predicted absolute binding affinities, and also capture the dynamic, atomistic details of the conformational changes induced by BH3 helices. Together, these details highlight the structural principles of BH3 in-groove engagement that determine apoptotic function, paving the way towards the modulation of the interactions among the Bcl-2 family members.

---

Programmed cell death pathways such as apoptosis require robust regulation to avoid malfunction. In the mitochondrial (intrinsic) pathway of apoptosis, regulation by the Bcl-2 family of proteins ensures precise commitment to cell death. The members of this family inhibit (e.g. Bcl-xL) or activate (e.g. Bid) apoptosis by regulating the perforation of the mitochondrial outer membrane (MOM) caused by effectors (e.g. by Bax) (1, 2). Alterations of the protein-protein interactions (PPIs) directly affect the regulation of apoptosis resulting in disease and disorders (3–5). The clinical success of other PPI modulators (6–8) underscores the therapeutic potential of targeting apoptotic PPIs. Crucial to the successful targeting of Bcl-2 protein family PPIs is the understanding of their diverse context consisting of distinct thermodynamic profiles, influences by the local environment (e.g. cytosol or membrane), and inherent conformational plasticity of the interaction partners. This multifaceted context together with the high degree of similarity between the protein members, and the overall network complexity of the Bcl-2 family make PPIs both promising and challenging targets for drug development (8–11).

Such Bcl-2 PPI regulators are already used within current leukemia therapies, since overexpression of apoptotic inhibitors (like Bcl-xL) suppress apoptotic activators (e.g. Bid) and effectors at the MOM as a primary driver of the disease. Current leukaemia therapies thus try to arrest tumour growth through inhibition of apoptotic inhibitors (8, 12), to restore apoptosis. Notably, tumor growth can recur if apoptotic effectors (e.g. Bax) acquire mutations which in turn change the PPIs to block the MOM perforating activity (13–15).

The conserved interface between the Bcl-2 homology (BH)3 domain and a hydrophobic groove in the Bcl-2 proteins mediates most PPIs within the Bcl-2 family network. However, while all family members contain an amphipathic BH3 domain, binding grooves are only present in effectors and inhibitors (2) (see Fig. 1B). In cytosolic conditions these grooves are usually occupied by the respective protein’s transmembrane C-terminal (*α*9) helix. Therefore, the groove becomes available for interactions with BH3 domains only upon membrane anchoring (16), through *in vitro* induced conformational changes (17, 18), or protein truncations (*α*9 deletion) (19, 20) (see Fig. 1A). The relative binding affinities of these conserved BH3 domain/groove interfaces determine how the Bcl-2 family network is regulated. BH3 helices have weaker affinities to inhibitors and effectors than their respective *α*9 helices and bind the hydrophobic grooves of inhibitors stronger than those of effectors (1, 2, 16, 17). This binding induces pro-apoptotic conformational changes in effectors leading to MOM perforation and apoptosis.

**Fig. 1.**
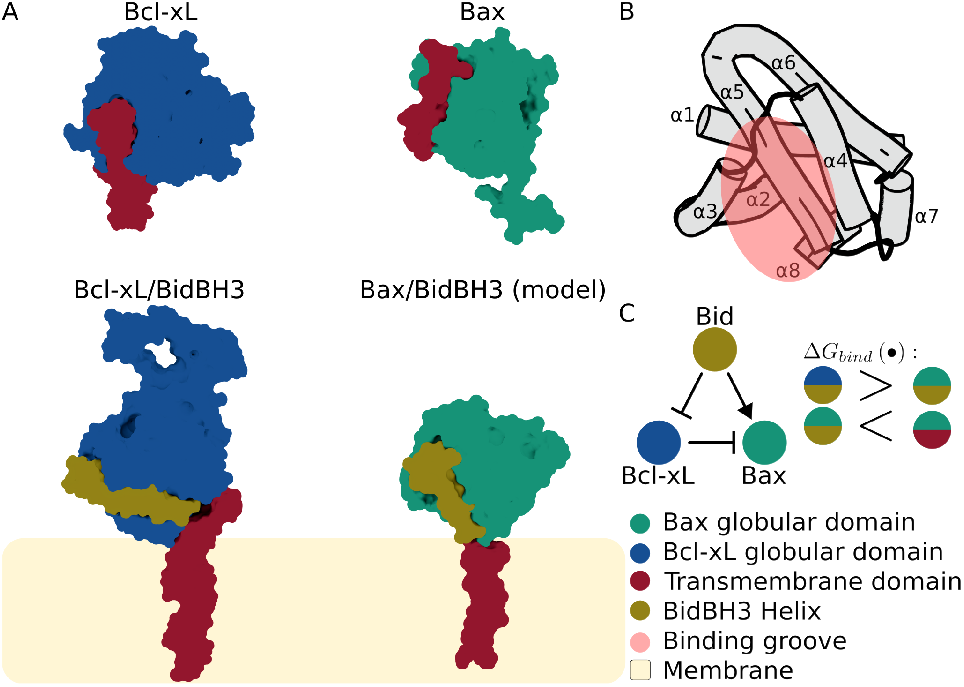
Structures and models of the interactions among BH3 domains, C-terminal transmembrane helices and canonical binding grooves. (A) Bcl-xL (blue) is an anti-apoptotic inhibitor partitioning between the cytosol and the mitochondrial membrane. In its cytosolic form, the transmembrane domain (*α*9 helix, red) is tightly bound to the groove; when inserted in the membrane, a Bid peptide (dark yellow) binds the available groove (12, 22). Bax (green) is a pro-apoptotic effector which is mostly cytosolic, but can reversibly translocate to the membrane. In its cytosolic form, the transmembrane domain (*α*9 helix, red) is tightly bound to the groove (PDB 1F16 (23)). In the putative membrane-anchored form, a Bid peptide (dark yellow) could bind to the available groove (17, 18, 23). Proteins shown as stylized surfaces. (B) Canonical conserved fold of Bcl-xL’s and Bax’s globular domain without the C-terminal *α*9 helix and *α*1*/*2 loop. Helices are shown as cylinders and the canonical binding groove is indicated with a red oval. (C) Scheme of a minimal interaction network of Bcl-xL, Bax, and Bid. Literature based binding affinity ranking associated with this minimal network indicated on the right.

Resolving the structure of a Bcl-2 family BH3 in-groove PPI gives detailed insights into the spatial arrangement of amino acids involved in interaction, but lacks information on the mechanisms and thermodynamics of PPI formation and dissociation of the complex. While kinetic and thermodynamic properties underlying PPIs can be experimentally obtained by e.g. surface plasmon resonance or isothermal titration calorimetry (21, 22), these methods lack atomic resolution. Linking kinetics and thermodynamics to structures of Bcl-2 family PPIs remains challenging due to this methodological gap and the general challenge of obtaining high resolution structures of the complexes, as full-length Bcl-2 complexes are often small, flexible, short-lived species occurring predominantly at the membrane.

Computational biophysical modeling can bridge this missing link between atomistic structures and thermodynamic properties (24–30). Structural modeling can resolve experimentally inaccessible states and PPIs, as it was recently shown for the full-length membrane anchored Bcl-xL bound to tBid (12). This model establishes a structural framework for investigating the interaction of a BH3 helix with the full-length, membrane-bound Bcl-xL and for extending this analysis to comparisons with membrane-anchored effector proteins such as Bax. Bax activation requires a MOM associated non-homomeric state (31, 32) which can be targeted by the BH3 domain of activators at the binding groove or at the non-canonical activation site (32, 33) and/or by the BH3 domain of other active Bax monomers (2, 15, 16, 34). Interaction with other proteins (18, 35–38) or lipids (39) inhibit or activate this MOM associated monomeric Bax (see Fig. 1). Starting from the structural homology with Bcl-xL and from available partial structures of Bax and BH3 helices (18, 40) we built a membrane-anchored monomeric Bax model with the transmembrane *α*9 helix inserted in the bilayer, with the groove available for BH3 domain engagement. Direct comparison of Bax in such an anti-apoptotic like, membrane-anchored form, with the membrane-anchored Bcl-xL capable of binding the BH3 domain of Bid at the membrane could provide insights into the structural determinants differentiating the fate of apoptotic inhibitors from effectors upon BH3 in-groove dimerisation.

Fundamental to such comparisons are the intra-molecular rearrangements occurring in Bax and Bcl-xL upon BH3 binding that could modulate the dynamics of the events (2, 41–43) and potentially propagate large conformation changes (44, 45). Differences in the PPIs could explain why interaction with apoptotic inhibitors lead to stable complexes (Bcl-xL/tBid), while complexes with effectors (Bax/tBid) are transient and might induce large conformational changes in a hit-and-run like manner (46, 47). While molecular dynamics (MD) simulations can investigate these intra-molecular dynamics they struggle to reach the timescales required for the observation of PPI association/dissociation. Collective variable (CV) based enhanced sampling methods allow accessing these events in atomistic MD simulations. Despite well-established CVs for the binding of small molecules to proteins in literature (48–50), the transfer of such CVs to the more ambiguous and expansive conformational landscapes of PPIs remains limited.

Here we probe the differences between the binding of BidBH3 helices to Bcl-xL and Bax to identify structural fingerprints of BH3 engagement that differentiate apoptotic function (see Fig. 1) (1, 2). To capture these elusive, transient events at an atomistic scale, we employed an advanced enhanced sampling molecular dynamics protocol (OneOPES paired with funnel restraints) capable of accelerating slow biomolecular binding timescales without losing atomistic accuracy (50, 51). This allowed sampling of the binding process and recovered Δ*G*_*bind*_ ranking under consideration of co-folding effects through combination with folding simulations of the BidBH3 peptide helix. Simulations revealed alternative bound states with differing pose selectivity, yet a conserved intra-receptor response to BH3 peptide engagement. These conserved interactions indicate that apoptotic function and BH3-triggered activation arise from differences in binding-groove rigidity, the *α*1*/*2 loop, and membrane-modulated BH3 pose selectivity and ambiguity. Together, these structural elements suggest why conversion of inhibitors into effectors (52, 53) is possible, whereas conversion of Bax into an inhibitor has not yet been shown.

## Energy landscapes of binding events to Bcl-xL and Bax

Investigating ligand-receptor binding and corresponding energetics in general requires accessing slow processes within the MD simulations. Here we ran funnel-OneOPES simulations with PPI-tailored CV’s to simulate the binding processes (see methods), thoroughly sampling along the binding events of investigated systems. To assess the validity of the method in recovering the available experimental affinity ladder, we computed the binding free energies (Δ*G*_*bind*_) for the two membrane systems of Bcl-xL/BidBH3 peptide and Bax/BidBH3 peptide, as well as a control system of Bax’s *α*9 to Bax itself. For each system we estimated the free energy surface (FES) along the distance between ligand and binding groove and calculated the Δ*G*_*bind*_ from the independent replicates (see Fig. 2). We then report the average across replicates of FES and Δ*G*_*bind*_ values. Estimates of the Δ*G*_*bind*_ stabilise after ~ 400 − 700 *ns* of simulation, when discarding the initial out-of-equilibrium fraction of the trajectories (100 − 150 *ns*) from estimation (see Fig. S4).

**Fig. 2.**
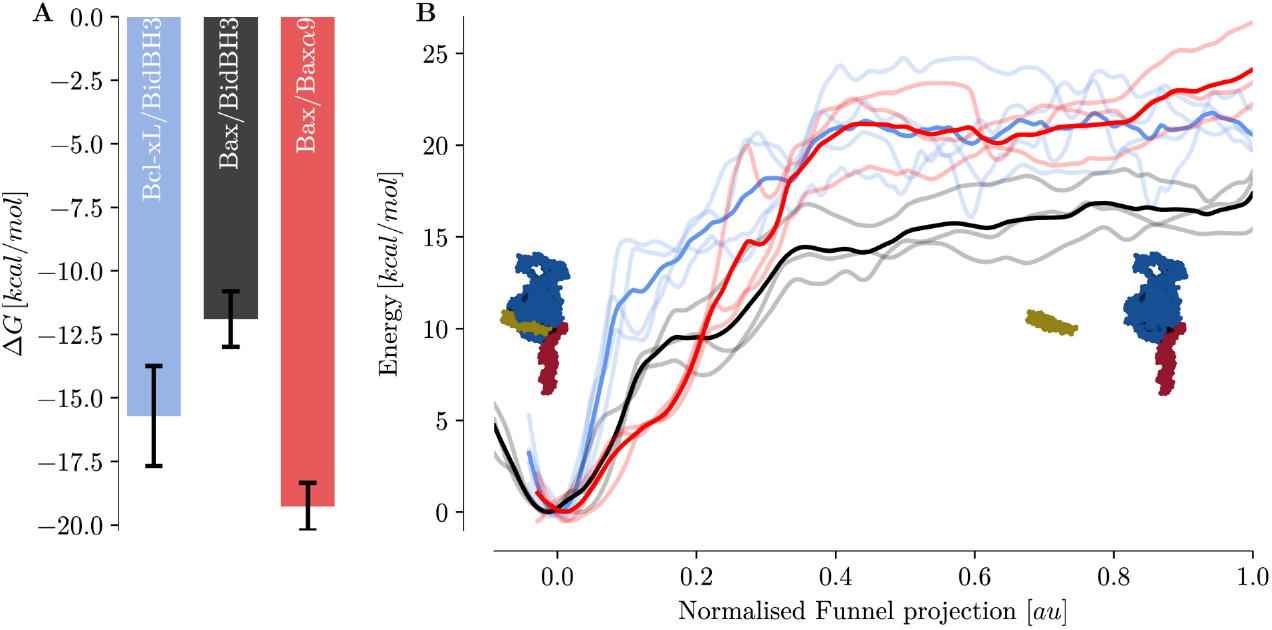
The Bcl-xL canonical binding groove binds BidBH3 peptide stronger than Bax. (A) Barplot of estimated Δ*G*_*bind*_ of the different systems. (B) FES of the simulated systems along the normalised distance between ligand and receptor (funnel projection). Independent simulation triplicates shown as transparent lines, average FESs shown as thick lines and standard deviation shown as transparent filled regions. Bcl-xL/BidBH3 peptide (blue); Bax/*α*9 (red); Bax/BidBH3 peptide (black).

Our calculations nicely reproduce the expected affinity ranking (see Fig. 2, Tab. 1) (1, 44, 54, 55). Notably, the re-folding event within the Bcl-xL *α*3 helix is pivotal to estimation Δ*G*_*bind*_ values for the Bcl-xL/BidBH3 peptide complex (see SI results, Fig. S1-S3). Initial estimates not taking the re-folding events within the *α*3 of Bcl-xL into account strongly overestimated the values (see Fig. S1, 2).

**Table 1.** Comparison between computed (comp) and experimental (exp) apparent binding energies of Bcl-xL and Bax to ligand helices. FL implies full length; mem. implies membrane; solv. implies solvent; Δ*C* implies C-terminal, mostly *α*9 truncation, Δ*L* implies replacement/shortening of *α*1*/*2 loop. The BidBH3 peptide_*f*_ helix is experimentally stabilised in the folded state by stapling (56). All energies given in [*kcal/mol*].

| | System | $\Delta G_{bind}^{exp}$ | $\Delta G_{bind}^{comp}$ | $\Delta G_{bind}^{comp} + \Delta G_{BH3-fold}^{comp}$ |
| --- | --- | --- | --- | --- |
| Bcl-xL | FL/BidBH3 pep. (mem.) | $-9.56 \pm 0.07$ (22) | | $-9.8 \pm 3.0$ |
| | $\Delta C$/BidBH3 pep. (solv.) | $-9.69 \pm 0.08$ (22) | | |
| | $\Delta C\Delta L$/BidBH3 pep.<sub>f</sub> (solv.) | $-9.89 \pm 0.16$ (56) | | |
| | FL/tBid (mem.) | $\sim -12.1$ (1, 44) | $-15.7 \pm 2.0$ | |
| Bax | FL/BidBH3 pep. (mem.) | N/A | | $-5.9 \pm 2.5$ |
| | FL/tBid (mem.) | $-10.92 \sim -10.62$ (1, 54, 55) | $-11.9 \pm 1.1$ | |

The estimated Δ*G*_*bind*_ between the BidBH3 peptide and Bax is in agreement with literature values between Bax and full-length tBid within the standard error of the estimate (see Tab. 1) (1, 54, 55). To strengthen the comparison with literature data, we also provide the estimate of the Δ*G*_*bind*_ between the Bax *α*9 and the canonical binding groove of Bax (Δ*G*_*bind*_ = − 19.28 ± 0.94 *kcal/mol*). This value is in agreement with the literature data, which consistently rank the Bax *α*9 helix as a stronger binder of Bax than the BidBH3 peptide (see Fig. 2) (1, 2).

## Correcting for BidBH3 peptide folding

Protein-peptide binding can induce fold changes in both binding partners (57, 58), but our binding protocol requires fold-restraints on the ligand helix to avoid conformational distortions upon contact with the funnel. BH3 peptides of apoptotic activators can be disordered before binding and fold upon/during binding to their interaction partners (59). An example of this is the BimBH3 peptide, which upon binding to truncated Bcl-xL in solvent gains a helical fold (60). Since both Bim and Bid are apoptotic activators, similar fold changes could occur. We therefore assessed whether restraining the BidBH3 peptide fold affects the binding event. Indeed, preliminary plain MD simulations of the BidBH3 peptide showed rapid unfolding of the helix in TIP3P water (see Fig. S5).

To gauge the energetic effect of the restraint, we estimated the folding component 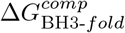 by simulating the folding of the BidBH3 helix in solvent (see Fig. S6). To achieve this, we employed a OneOPES protocol by Rizzi *et al*. (61) calculating the 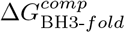 of the BidBH3 peptide from three independent simulation replicates (see Fig. 3). This showed that the BidBH3 peptide helix is strongly disposed towards the unfolded state in solvent without a local minimum in the *α*-helical state, corroborating the preliminary plain MD simulations. We therefore incorporated the helix folding to correct the Δ*G*_*bind*_ estimates via:

**Fig. 3.**
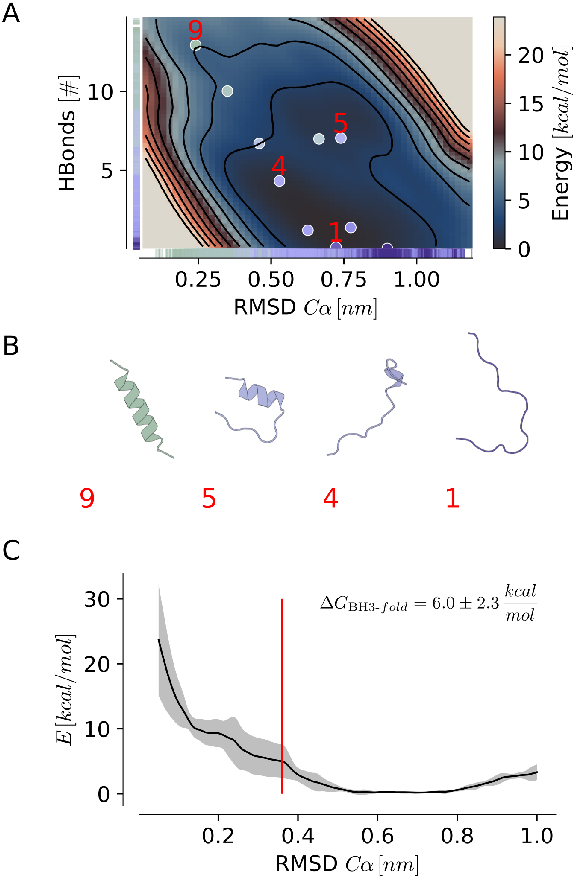
The Bid BH3 helix is unstable in solvent. (A) Two dimensional FES associated with BH3 helix folding expressed in the RMSD_*Cα*_ and hydrogen bonds (HBonds) space, with RMSD_*Cα*_ calculated with respect to BH3’s folded state and *α*-helical hydrogen bonds measured between the backbone atoms. Representatives of clusters within the structural ensemble indicated as dots within the FES. (B) Structural representatives (cartoon) along the folding process extracted from (A) (cluster indicated in red). (C) Folding free energy calculated along the RMSD_*Cα*_ between the folded state and unfolded states based on three independent simulation replicas.

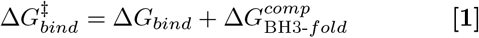

where we decouple the process into its binding and folding components (see Tab. 1).

Including this correction of the observed Δ*G*_*bind*_ for Bcl-xL/BidBH3 peptide results in a 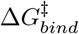 in line with the experimental values. For the Bax/BidBH3 peptide system the correction leads to underestimation of experimental Bax/tBid values (see Tab. 1).

## Binding groove closure upon BH3 binding correlates with distant sites

The canonical binding grooves of Bax and Bcl-xL have similar architectures, where the groove itself and the *α*3 helix represent a hydrophobic surface and the *α*4 helix and the *α*4*/*5 loop represent a dipolar electrostatic surface. Bid’s BH3 helix is amphipathic, aligning with the hydrophobic surface into the groove and orienting its charged dipolar surface complementarily to the receptor (see: Fig. 4A). Instead, when receptors are unbound with their transmembrane *α*9 helix inserted in a membrane, water exclusion from the hydrophobic patches of the binding groove require conformational changes within the groove.

**Fig. 4.**
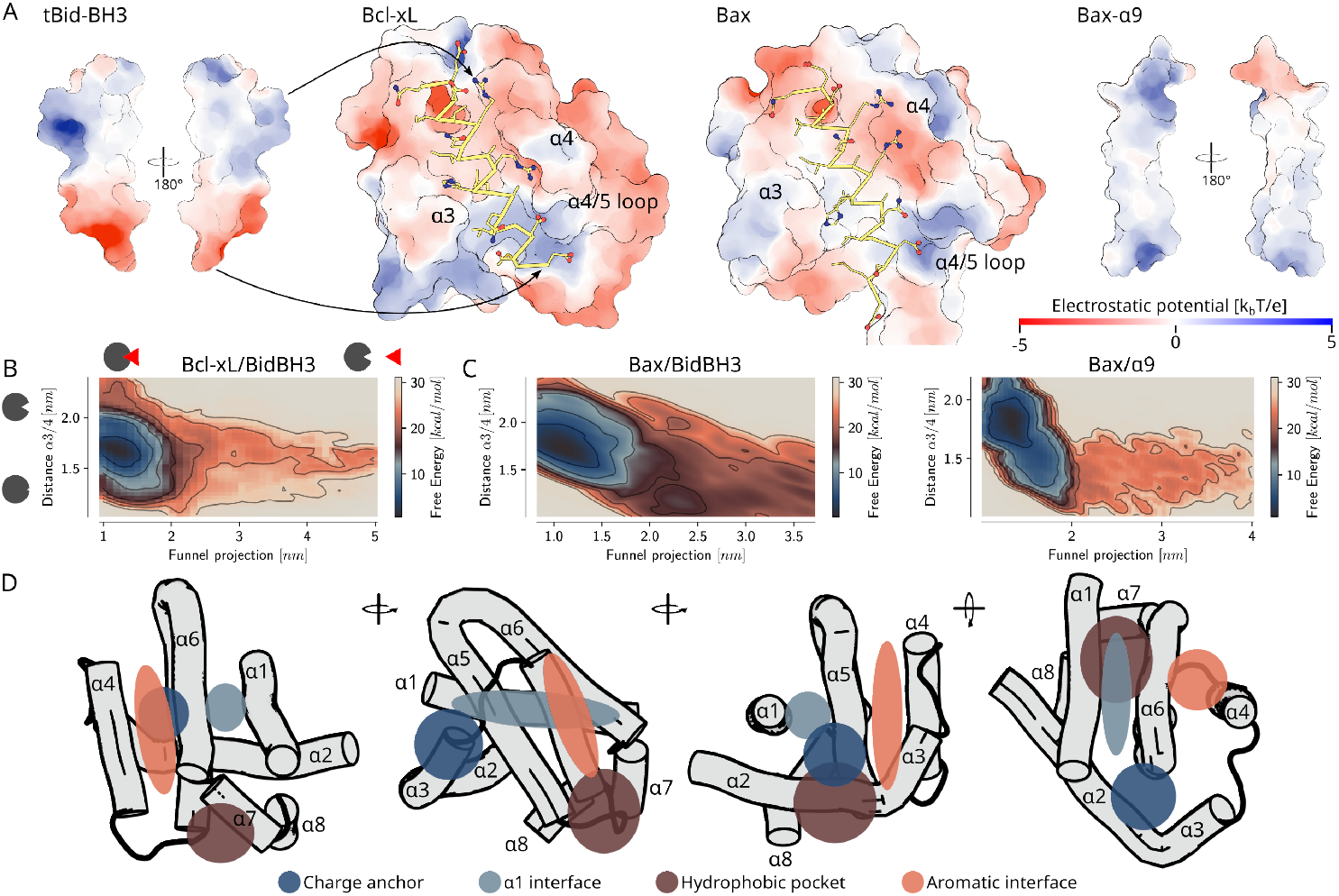
The canonical binding groove of Bax quickly closes in unbound states. (A) Electrostatic potential calculated through the linearised Poisson-Boltzmann equation mapped onto the surface of BidBH3 peptide, Bcl-xL, Bax, and Bax’s *α*9 helix. Ligand helices in Bcl-xL and Bax indicated as yellow ribbon with sidechains shown as lines and spheres coloured by element (blue: nitrogen; red: oxygen; yellow: carbon). Ligand helix electrostatic surfaces oriented in relation to the binding grooves in Bcl-xL and Bax. (B) average FES across independent simulation replicas of the distance between *α*3 and *α*4 against the funnel projection for the Bcl-xL/BidBH3 peptide system. Binding groove closage and binding event along the two dimensional FES indicated as pictograms. (C) average FES across independent simulation replicas of the distance between *α*3 and *α*4 against the funnel projection for the two Bax systems. (D) Schematic representation of regions within the canonical globular Bcl-xL/Bax fold that contain correlated changes with the binding and groove closage event. Individual correlations of distances are in supplementary figures S11 to S12.

Indeed, both receptors display conformational changes of the binding groove during the binding process. Upon binding the binding groove of Bax opens through increase of the distance between the *α*3 and *α*4 helices (see Fig. 4C) in both the membrane associated Bax/BidBH3 peptide and solvent Bax/*α*9 binding simulations. In contrast, Bcl-xL’s binding groove only shows correlation between binding of the BH3 peptide and the *α*3*/α*4 distance (see Fig. 4B) when the *α*3 helix of Bcl-xL gains fold (see Fig. S7 and Fig. S8). This fold gain in the Bcl-xL *α*3 helix correlates with decreased binding groove accessibility, with the bound BidBH3 peptide stabilizing the helices unfolded state (see Fig. S2D).

To investigate intra-receptor responses to the binding event, we selected inter-residue distances within the receptor proteins (Bax or Bcl-xL) that show high correlation with the binding event (funnel distances) and calculate their correlation with further macroscopic features such as groove accessibility and ligand helix orientation. Distances identified by this selection predominantly showed the strongest correlations with the groove accessibility, suggesting that changes within the receptor result from groove closure and not the binding event directly. In general, membrane associated Bax and Bcl-xL showed similar regions of interest (see Fig. 4D), and correlating distances qualitatively agreed between the Bax/BidBH3 peptide simulation and the Bax/*α*9 control system in solvent (see Fig. S11 to S13). Markedly, some specific interactions identified within these regions are conserved between Bax and Bcl-xL (residues involved are listed in Tab. 2).

**Table 2.** Binding and groove closure correlate with conserved residues in the globular fold. Table lists binding sensitive residues and their location in the globular fold (see Fig. 4D) observed in both membrane associated Bax and Bcl-xL, for which experimental evidence exists in either Bax or Bcl-xL.

| Region | Bax Residues | Bcl-xL Residues | Effect | Literature context |
| --- | --- | --- | --- | --- |
| Aromatic interface | R109 | R139 | Interacts with the ligand helix in bound state, but interacts with the α5/6 loop in the unbound state | Charge reversal mutation in Bax results in a loss-of-function (40) |
|  | W139 | W169 | Connects the α4 helix to the α1 helix and α1/2 loop through the α5/6 hairpin (Bax<sup>I31</sup>, Bcl-xL<sup>L13</sup>, see Fig. 4D, lightblue) | Mutation in Bax disrupts allosteric signalling (32); vMIA (viral mitochondria-localized inhibitor of apoptosis) exploits this connection to inhibit membrane associated Bax (35, 36). |
| Charge Anchor | R37, D68 | K16, D97 | Interaction lost in active conformations (40, 62, 63); anchors the α1 helix to the α2 helix on the backside of the α3 helix (close to Bax: D71, Y115, K119) | Both Bax residues were also identified in previous simulation studies (11, 45); Bax<sup>R37</sup> is part of an epitope (18, 38); Charge reversal mutation Bax<sup>D68R</sup> deregulates Bax (64). |
| Hydrophobic pocket | I19, W158, W107 | N5, W189, W137 | Connects α1 helix to the binding groove through the bottom of the α5/6 hairpin and the α4 helix | In Bax, this membrane proximal hydrophobic interface harbours a pocket gated by W107 and I152 whose binding by lipid tails increases Bax activity (39). |
| Binding groove | F116 | F146 | Sensitive to groove closure in solvent Bax (Bax/α9 binding) and to fold gain in Bcl-xL α3 within membrane bound Bcl-xL | Identified in previous simulation studies as signature of crypticity in Bcl-xL (65) |

In summary, the binding of ligand helices (BH3 or helix *α*9) to Bax correlates with binding groove closure and changes in amino acid clusters within the protein’s core connecting the binding groove to the *α*_5*/*6_ helix latch and the inhibitory *α*1 helix through both hydrophobic and charged interfaces. Membrane bound full-length Bcl-xL shows homologous correlations and interactions, but (as previously shown in aqueous solution (60)) requires changes in the fold of *α*2 and *α*3 for binding groove closure. Paired with the longer *α*1*/*2 loop in Bcl-xL this difference might suppress family conserved intra-receptor signalling between the binding groove and *α*1 (BH4) helix.

## Pose ambiguity differentiates Bax from Bcl-xL

Despite the complementary electrostatic dipoles of both the BidBH3 peptide helix and the receptor binding groove the BidBH3 peptide helix binds both Bax and Bcl-xL in two orientations (see Fig. 4, C): The canonical orientation in which the C-terminal of the ligand helix binds to the membrane proximal sub-pocket, and the flipped orientation in which the C-terminal binds to the (cryptic (60, 65)) membrane distal sub-pocket (see Fig. 5, A). This flipped bound state is predominantly stabilized by hydrophobic interactions, misaligning the electrostatic dipoles of the ligand helix and the receptor’s binding groove.

**Fig. 5.**
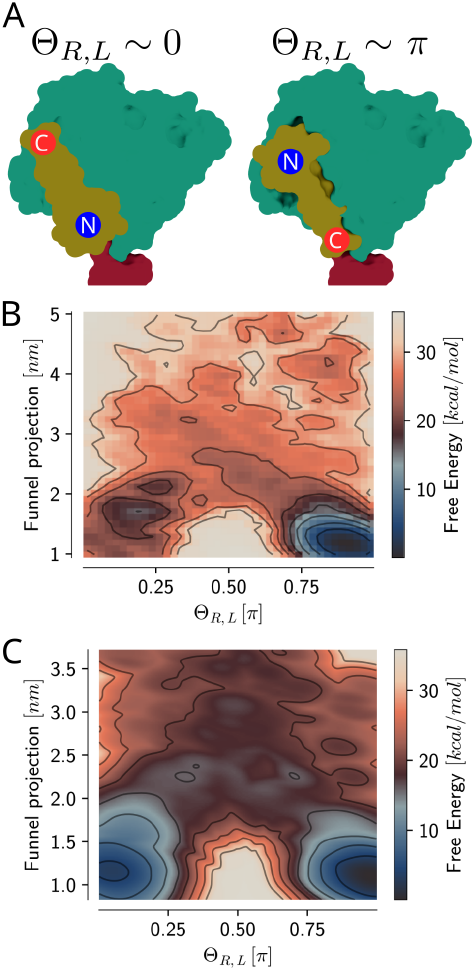
Opposite orientations of the BidBH3 peptide bound to Bcl-xL and Bax. (A) Simplified surface representations showing the two possible bound states: flipped (left) and canonical (right). (B, C) FES’s show the potential along the funnel projection and ligand orientation relative to the receptor. (B) FES of the Bcl-xL/BidBH3 peptide system. (C) FES of the Bax/BidBH3 peptide system.

While our simulations show that both Bax and Bcl-xL present these two opposed binding poses, Bax’s binding groove shows a hint for stronger pose ambiguity than Bcl-xL. In Bax, the difference in free energy between the flipped and canonical bound state 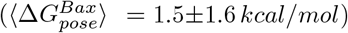 is smaller than in Bcl-xL 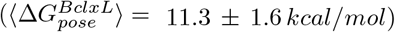 (see Fig. 5). This might be a consequence of the weaker electrostatic dipole and stronger hydrophobicity within Bax’s binding groove compared to Bcl-xL as well as the weaker correlation of intra-receptor distances with the ligand orientation in Bax (see Fig. 4, S11, S12). The identified stronger pose ambiguity within Bax compared to Bcl-xL might provide a key for the different fate of the effectors and inhibitors upon BH3 binding.

## Discussion

The different protein-protein affinities within the Bcl-2 family protein interaction network play a key role in regulating intrinsic apoptosis.

Therefore, advancing our understanding of biology and developing new drugs that target intrinsic apoptosis requires an understanding of the atomistic details that determine these affinities. To this end, we employed enhanced sampling MD simulations to explore the binding mechanisms and the energy landscapes associated with BidBH3 peptide binding to full-length membrane bound Bcl-xL and Bax.

Our approach yielded results consistent with the available experimental data, providing insights into the dynamics of the binding groove and ligand helix co-folding, as well as the role of membranes during binding. Still, it is worth stressing that both computational and experimental estimates of binding affinity (Δ*G*_*bind*_) are subject to limitations. The former are affected by the accuracy of force fields and sampling, while the latter are affected by varying experimental conditions and the use of different constructs (e.g. tBid vs the truncated BidBH3 peptide or solution vs membrane)(66). Although these factors hinder exact quantitative comparisons, our results remain consistent with existing experimental data. Indeed, the total (binding + folding) Δ*G*_*bind*_ of the BidBH3 peptide to Bcl-xL agrees with the experimental binding affinities obtained in similar conditions (1, 22, 44). At the same time, our Δ*G*_*bind*_ estimates for Bax is in line with experimental values for the full-length tBid (1, 54, 55) (see Tab. 1). In this case however, since in tBid the helix is folded (67), the closest comparison is with the computed binding free energy. What is more, our relative ranking agrees with available evidences. This implies that our protocol and our models (see Fig. 2 and Tab. 1) are able to capture the main differences between the systems.

The agreement between the experimental and computed free energies allows us to analyse in detail the mechanistic and energetic differences between Bax and Bcl-xL. First, the binding groove of Bax is less accessible than the one of Bcl-xL. This is due to the fact that while Bax’s *α*3-helix stays folded and can promptly close the groove, in the case of Bcl-xL it has to refold before closing the groove (see Fig. 4B, Fig. S1, Fig. S8). This is in qualitative agreement with previously reported behaviour in solution (60, 65, 68). Yet, our simulations show that the membrane shifts the equilibrium towards greater binding site accessibility through stabilization of the unfolded Bcl-xL *α*3 helix (see Fig. S2). This observation is in line with apo Bcl-xL membrane simulations and nuclear magnetic resonance spectroscopy (NMR) data with unfolded flexible *α*3 helices (12, 22).

Second, the binding modes of the Bax/BidBH3 and Bcl-xL/BidBH3 heterocomplexes are different. In the case of Bax, the BidBH3 peptide helix binds in both the canonical and flipped pose, while in Bcl-xL it preferentially binds in the canonical pose (see Fig. 5). This geometry arises from the stronger dipole moment of the Bcl-xL groove and contributes to the differential binding affinity. In light of Bax’s active in-groove dimerisation resembling a flipped binding pose (40) and Bcl-xL’s ability to similarly bind the BaxBH3 helix (69), pose ambiguity appears crucial to apoptotic effector function.

Despite these two main discrepancies between Bax and Bcl-xL, we also observed key similarities in how the two proteins respond to binding and groove closure (see Fig. 4). For Bax, the allosteric connection from the binding groove to the inhibitory *α*1 helix on the proteins backside is well established (11, 16, 32, 45, 70). Similar to Bax, if Bcl-xL loses its inhibitory *α*1 helix (through mutation or cleavage (71)), Bcl-xL perforates the MOM. This mechanism is used during mitosis regulation to induce intrinsic apoptosis (53). The Bax homologous residue-residue interactions identified within this study show that within Bcl-xL binding groove and *α*1 helix are similarly linked.

In solution, BH3 peptides typically exhibit an unfolded state and must co-fold upon binding to their respective receptors. Previous experiments have shown that stabilizing the peptide’s *α*-helical content (through “stapling”) enhances Bax activity and binding kinetics by reducing the entropic penalty, though it also introduces compensating enthalpic changes that complicate direct comparisons with our simulated fold-restraints (22, 56, 60, 72). However, in a physiological context, the membrane environment itself likely acts as a natural stabilizer for the amphipathic BidBH3 helix (73). Our unbiased simulations support this, demonstrating increased helical stability when the peptide is membrane-associated (see Fig. S5). Given that Bcl-xL increases the membrane association of tBid (74), a simple “folding-upon-binding” mechanism may not fully capture hetero-dimerization *in vivo*. Instead, complex formation likely involves a spectrum of mechanisms dependent on the receptor protein, ranging from coupled folding-and-binding in the cytosol to the binding of pre-folded, membrane-bound helices.

Beyond its stabilization effects, the membrane also physically restricts the available binding modes (see Fig. 5) by anchoring the full-length activator proteins. For instance, because the BH3 helix in full-length tBid is anchored to the membrane at its C-terminus, the C-termini cannot easily reach the membrane-distal sub-pocket of the receptor (12). This physical tethering likely biases the interaction toward the canonical binding pose, whereas an N-terminal anchorage might favour the flipped pose (72, 73). Clarifying how the membrane guides both ligand helix stabilization and binding-mode selection provides vital insights into how protein-protein interactions are regulated across the Bcl-2 family.

In conclusion, our enhanced sampling simulations of membrane-anchored Bax and Bcl-xL reveal critical structural determinants that dictate their distinct apoptotic functions. Despite showing similar intra-protein signalling upon BH3 peptide binding, Bax is fundamentally different from Bcl-xL due to its higher binding-groove rigidity, greater pose ambiguity, and a shorter *α*1*/*2 loop. Furthermore, the membrane plays a crucial, active role in tuning the affinities of these hetero-complexes by promoting BH3 peptide folding and spatially restricting binding orientations. Together, these differential structural and dynamical features provide a clearer mechanistic picture of apoptotic regulation and offer targeted pathways for the future development of conformation-specific therapeutics.

## Materials and Methods

### Structure preparation and models

An apo monomeric membrane associated structure of human Bax was generated via homology modelling using MODELLER (75). The homology model was based on the catfish Bax PDBID: 5w63 (18) and its globular state PDBID: 1f16 (23). Models were generated using the AutoModel functionality without additional restraints. The slow autoscheduler and MD refinement were used with a maximum of 6000 conjugate gradient repeats and eight optimization repeats. Models were discarded if their molpdf score was above 1 × 10^6^. Generated models were re-scored to minimize the sum of their molpdf and DOPE-HR score. The highest ranking model was used as apo structure.

The holo membrane associated Bax-monomer was generated from the apo-structure. A BidBH3 peptide was superimposed onto the structure from PDBID 4zig (17) and termini methylated with PyMOL (76) (ACE, NME). Side-chain clashes were removed via re-docking with Haddock (77). During Haddock re-docking surface contacts were turned on and ambiguous restraints off. 200 rigid body dockings were performed and minimized ten times and 200 structures were semi-flexibly refined and analysed. Within the docking, no random starting orientations were used, the translations were turned off, the cross-docking disallowed, and no initial docking performed. No MD refinement was conducted. The resulting structures contained only one cluster and its representative was used as starting frame in simulations. The Bcl-xL/BidBH3 helix complex was extracted from the Bcl-xL/tBid complex previously published (12) by reducing tBid to residues Q79 to D98 and termini methylated with CHARMM-GUI (78) (ACE, CT1).

For the solvent Bax simulations the PDBID 1f16 NMR structures was used (23). Folding simulations of the BidBH3 peptide were based on the termini methylated ligand helix extracted from the Bcl-xL/BidBH3 peptide complex described above.

### Simulation box setup and parameters

Solvent states and apo membrane states were protonated with pdb2pqr and propka to a pH of 7.5 (79, 80). Membrane associated holo models and the apo BidBH3 peptide helix were protonated with CHARMM-GUI to a pH of 7.5 (78). For protein charges see Tab. S1.

Membrane associated systems were built and solvated with CHARMMGUI (78) in a membrane of 9,10 and 12,13 unsaturated cardiolipin (TLCL) (20 %) and 1-palmitoyl-2-oleoyl-sn-*glycero*-3-phosphocholine (POPC) (80 %) in a hexagonal box. Lipids were equally distributed across leaflets. Solvent systems were solvated with GROMACS in a dodecahedron boxes. For the ligand helix box, box size was estimated through first simulating the helix’s unfolding in vacuum by breaking all backbone hydrogen bonds with adiabatic biasing MD (81, 82). All boxes were charge neutralised with sodium chloride and sodium chloride added to a 150 *mM* concentration. For box compositions see Tab. S2. Boxes were parameterised with the CHARMM36m force field and corresponding CHARMM36m TIP3P water (83).

All simulations were conducted with GROMACS 2023 and PLUMED 2.9.0 (82). Constant simulation parameters are summarized in the following, while changing parameters during the equilibration are listed in Tab. S3. Production simulations were run with a time step of 2 *fs* in the NPT ensemble. Temperature was controlled with the V-rescale thermostat (310.15 *K*; *τ* = 1 *ps*) (84), separately coupling membranes (if present), proteins and solvent. Pressure was controlled with the C-rescale barostat (1 *atm*; compressibility = 4.5 × 10^−5^ *bar*^−1^; *τ* = 5 *ps*) (82) with isotropic or semi-isotropic coupling depending on whether a membrane was present in the box or not. Lennard-Jones interactions were treated with the Cut-off scheme using force switching at a neighbour list cut-off of 1.2 *nm* and switch radius of 1.0 *nm*. Coulomb interactions were calculated using PME with a cut-off 1.2 *nm*. Neighbour lists were calculated with the Verlet cut-off scheme. Hydrogen bonds were restrained with LINCS (85) in all simulations and center of mass motion removed every 100 steps on membrane, protein and solvent groups. Frames were written every 100 *ps* in binding simulations and every 10 *ps* in folding simulations.

### Protein Protein binding simulations

Contributions from electrostatic and van-der-Waals interactions, hydrophobic effects, and conformational rearrangements guide binding on different time and spacial scales in a case dependent manner (58, 86, 87). While individual interactions are often weak and unspecific, their synergies can form stronger global interactions collectively determining the bound state through alignment of hydrophobic patches and dipole moments. This is especially relevant for PPIs resulting from few different interaction types, but large interaction surfaces (57, 58, 86–88). Protocols sampling the binding of PPIs should consider these different scales and synergies while optimizing computational cost, maintaining structural integrity and capturing non-accelerated faster-than-binding degrees of freedom.

To sample the dissociation and association of the ligand BH3 helices from and to the receptor Bcl2-proteins biased enhanced sampling was conducted with OneOPES (50, 51) under a funnel restraint (48, 89). OneOPES is a replica exchange based biased enhanced sampling protocol constructing bias potentials on-the-fly with OPES (90, 91). This necessitates description of the binding process by collective variable(s).

Simulations were driven predominantly along two CVs accompanied by auxiliary CVs biased within the OneOpes scheme (see Fig. 6 and Tab. S4,S5), similar to those used for small molecular ligands (50). Binding processes between proteins are driven by interactions with different decays and reaches. Especially in protein/protein interactions dipole moments and orientations between the partners influence the binding process. CV’s aiming to describe this process thus should benefit from combining information along multiple ranges of interactions. The first main CV was the distance between ligand and receptor along the funnel 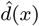 and the second a multi-level contact-map *S*(*x*) aiming to describe interactions from encounter to bound complexes with a non-zero gradient along the full (un)binding event. This contact map combines short-range (*SR*) residue-residue interactions, intermediate range (*IR*) interaction between subgroups of the ligand helix with the binding groove as well as long range (*LR*) interactions between ligand and receptor. This allows description of atomistic details during the early stages and coarse-grained details of the later stages of the unbinding process by extending the contactmaps range of non-zero gradient (see Fig. S14, A). Each contact (*SR, IR*, and *LR*) is described by a rational switching function:

**Fig. 6.**
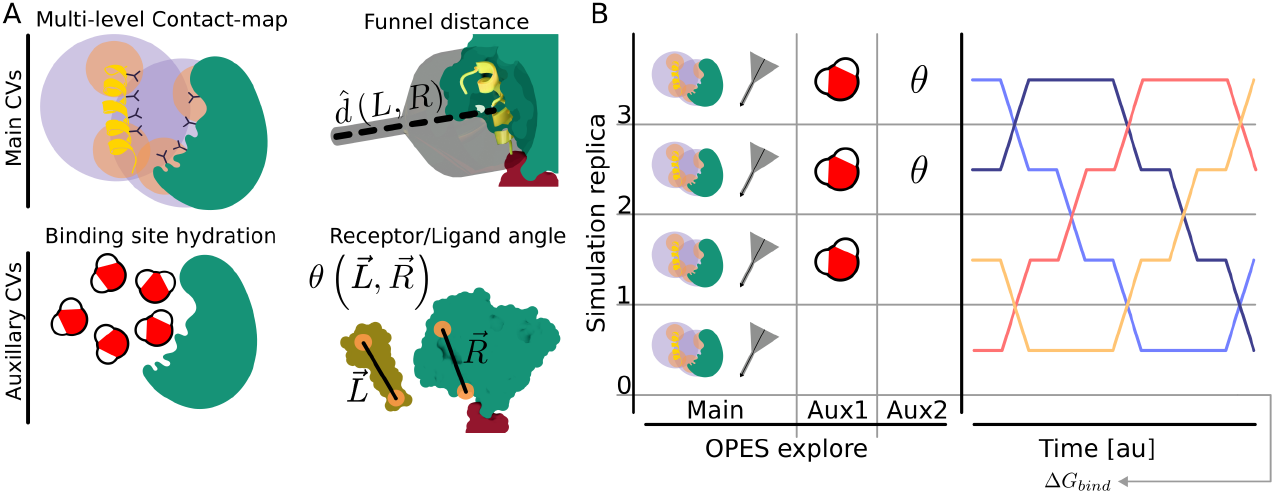
CVs utilized and simulation protocol. (A) Visual descriptions of cornerstone CVs used within the OneOPES protocol used to sample binding events. Receptor protein shown as stylized green surface and ligand as stylized orange cartoon or surface. Contact levels indicated as transparent spheres and residues (‘Y’). Funnel indicated as gray glass with dashed line as funnel vector. (B) Schematic overview of utilized protocol. Four replicates with either only the main CVs, or increasing amount of auxiliary variables (Binding site hydration; Ligand/Receptor angle) biased were simulated under replica exchange. FES and Δ*G*_*bind*_ values were extracted from the base replicate, only experiencing bias on the main CV’s (multi-level contactmap; Receptor/Ligand distance).

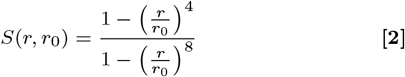

where *r* is the distance between atoms of the contact and *r*_0_ sets the reference distance of the contact. Within the contact-map *r*_0_ differs depending on the contact type (short: 0.8 *nm*, intermediate: 1.6 *nm*, long range: 3.2 *nm*). The different *r*_0_ were chosen such, that the gradient of the contactmap has no minima besides the global minima in the bound state and extends to non-zero values up to the unbound state (see Fig. S14, B). The value of the contactmap *S*(*x*) is then determined as the sum across all contacts:

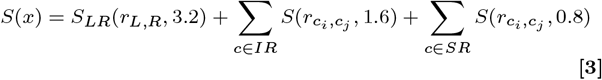

To accelerate sampling two relevant auxiliary CV spaces were biased: (a) the hydration of the binding groove, and (b) the angle between the binding groove and ligand helix (Θ_*R,L*_). This angle was described within a feature switching CV, that switches between the angle and the distance between ligand and receptor:

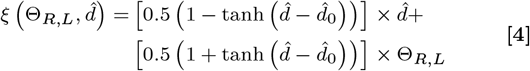

where 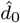 is a free parameter to position the switch along the funnel projection. We smoothly switch between biasing 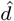 and Θ_*R,L*_ to accelerate rotation of the ligand in relation to the receptor only in unbound states and reduce unphysical biases on the receptor protein. This is a highly redundant CV by itself, but combination with the multi-level contact map *S* or funnel projection 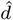 results in an informative two-dimensional CV-space used within auxiliary biases.

For the final Bcl-xL/BidBH3 peptide simulations the fold state of the *α*3 helix was also accelerated as auxiliary CV and the auxiliary CV’s reordered (see Tab. S5). This CV is defined as the sum of hydrogen bonds present in the fully folded state of the helix but not in the bound “un”-folded state of the helix:

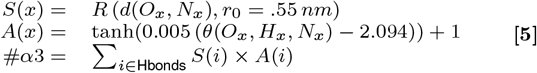

where [*O, H, N*]_*x*_ are the atoms involved in the corresponding backbone hydrogen bond, *R*( ·) is a rational switching function (see Eq. 2), and *θ*( ·) is the hydrogen bonds angle.

To improve local sampling additional restraints constrained the accessible conformational space. The positional space available to the unbound ligand was constrained with a sigmoid funnel restraint (89) with a *κ* of 20000 *kJ/mol* in respect to the ligand center of mass (CoM). To avoid conformational artifacts due to the funnel, backbone hydrogen bonds of the BH3 *α*-helices were restrained with lower walls slightly below their fully folded value (see Fig. S14, C and Eq. 5). This still allows for the unfolding of up to one *α*-helical loop and structural flexibility without full unfolding. These hydrogen bond restraints had a *κ* of 1000 *kJ/mol*. In the Bcl-xL/BidBH3 system and Bax systems some additional restraints were necessary and are listed in Tab. S7.

OneOPES simulations were conducted with the same simulation settings and parameters as described above. Deviating from the published OneOPES (51) protocol, here only four replicas are simulated in a replica exchange scheme and no multi-thermal bias used. This was done due to system sizes and to save on computational resources. Exchanges were attempted every 2000 steps. Re-weighting and analysis were conducted on replica zero of each simulation. For each biological system, three or four independent OneOPES simulations were run for 800 *ns* or 1400 *ns* each (see Fig. S15, S16, S17).

### Ligand helix folding

To account for the effect of ligand helix co-folding on the Δ*G*_*bind*_ the folding free energy (Δ*G*_*fold*_) of the BidBH3 peptide helix in solvent was estimated. Estimates of Δ*G*_*bind*_ were computed using a previously developed folding specific CV and associated OneOPES protocol (61). In brief, this CV describes a combination of both side chain contacts and hydrogen bonds of folded and unfolded states allowing distinction of the two states and their sampling (61). To identify which contacts and hydrogen bonds to consider within the CV initial simulations of the folded and unfolded state were conducted. As the folded state simulation showed rapid unfolding (see Fig. S5), instead contacts and hydrogen bonds in the folded state were identified from a restrained folded simulation. Three independent replicates of the folding OneOPES protocol (see Tab. S6) were run for ~ 700 *ns* and the first 100 *ns* discarded from analysis.

### FES calculations and reweighting

From all simulation trajectories only the base replica zero of the OneOPES scheme were analysed after discarding the first 100 *ns* of sampling. Simulation frames were re-weighted with the total bias applied to each frame, where weights are:

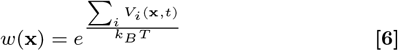

The computed weights were used in all FES calculations and otherwise re-weighted calculations based on the trajectories.

All presented 1D FESs were calculated with weighted Kernel density estimation (KDE) using the KDE implementation of scipy in python. Presented 2D FESs were calculated with a bandwidth of 0.1 (except the Bcl-xL/BidBH3 groove closage FES; 0.175) and without consideration of weights in the covariance matrix calculation. The energy values were then computed with:

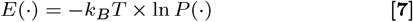

and the minima shifted to zero. For all plots values were extracted from this functional at a plot dependent spacing along the CV space.

FESs for the estimation of Δ*G*_*bind*_ were calculated along the funnel projection vector (*z*) with a bandwidth defined per system^∗^. Each of the independent replicates were processed separately and the final free energy projection calculated as mean across replicates. For each replicate the Δ*G*_*bind*_(*z*) was calculated by defining bound and unbound (solvent) states and taking the difference between the integral over the FES of each state. States were defined by segmentation of the re-weighted histogram along the funnel projection into three states via the Otsu algorithm. The bound state is defined as the range of 2 × *σ* around the mean of the lower projection state after the first otsu split. The unbound (solvent) state is defined as the range of 2 × *σ* around the mean of the highest projection state after the second otsu split. By automatically selecting the two states of the system through this procedure, estimation of the Δ*G*_*bind*_ is based on those regions of the FES that are based on the most sampling. This gives an indirect qualitative measure of sampling quality. As the unbound state of binding simulations is assumed to be quite diffusive, an unbound state that is defined by small range of values along the funnel projection is indicative of insufficient sampling.

Simulations experience a funnel restrain in the unbound states. Δ*G*_*bind*_(*z*) were funnel corrected for the restraint applied by the funnel with (48, 89):

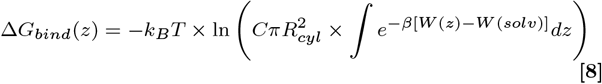

Where *C* is the standard concentration and *R*_*cyl*_ is the radius of the funnel in solvent. Funnel corrections remove the space dependence from the sampling (48, 89), correcting for the limited diffusion of ligand molecules and shifting the entropic contribution accordingly. As the *α*9-helix of Bax is covalently bound to the receptor (Bax) no funnel correction was applied in the calculation of Δ*G*_*bind*_ for this system.

FESs for the estimation of Δ*G*_*fold*_ were calculated along the C*α*-root mean squared deviation (RMSD) to the folded state of the ligand helix with a bandwidth of 0.075. Each independent replica was processed separately and the seperatrix between folded and unfolded states set to a value of 0.36 *nm*. No local minima in the folded state was present in the FESs complicating the clear definition of the folded state and seperatrix. To estimate the seperatrix we considered the fluctuation of the C*α*-RMSD of the BidBH3 peptide helix in folded unbiased membrane bound simulations, defining it as the 90 % quantile of C*α*-RMSD to a folded reference within that trajectory. The Δ*G*_*fold*_ is thus defined as:

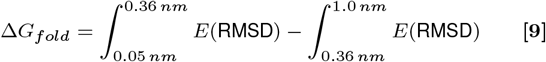

### Analysis

Additional features, such as hydrogen bonds, distances, and angles not monitored during simulation were computed with MDAnalysis and custom wrapper modules (92). If needed, such features were used to generate FESes (see FES calculations and reweighting). Hydrogen bonds were considered up to a distance cutoff of 3.5 *Å* between donor and acceptor atom and between 180 − 120^°^ angles. *α*-helical hydrogen bonds were defined as hydrogen bonds between backbone atoms that have a residue sequence separation of 4. *α*H-bonds were measured as distance between acceptor and donor atoms transformed with rational switching functions (*n* = 4; *m* = 14; *r*_0_ = 5.5 *Å*).

Dimensionality reduction of the all-atom space was achieved with GPU re-implementation of ELViM (93, 94) in python with numba. Features of interest and energies of individual frames in the dimensionality reduction were mapped in plots for initial visual feature analysis (see Fig. S8 to S10). States were extracted from the projection with Gaussian mixture model clustering after cluster number identification with OPTICS. Cluster representatives were extracted based on the dissimilarity matrix between intra cluster frames in a re-weighted manner.

We probed residue level information within simulations in relation to identified macroscopic features such as funnel projection (ligand/receptor distance), ligand helix orientation, contactmap, binding groove distance (*d* (*α*3, *α*4)), and (if present) Bcl-xL *α*3 helical fold. Distances within receptor proteins excluding N-terminal loops and trans membrane *α*9 helices were calculated between relevant sidechain heavy atoms. In the Bcl-xL/BidBH3 peptide system the *α*1*/*2 loop was also excluded from analysis due to its strong conformational flexibility and unlikeliness of exhaustive sampling in the unbound state. Considered sidechain heavy atoms were: Nitrogens and Oxygens for polar and charged amino acids, for ring containing amino acids except Proline all carbon-atoms except C*β*, and for Proline and unpolar amino acids all carbon-atoms. Using these selections both the pairwise distance and contact matrices were calculated where a contact was defined with following switching function:

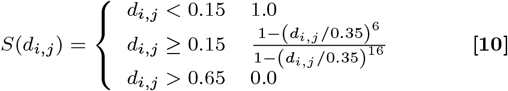

where all distances and cutoffs were in *nm*. Matrices were pre-filtered by removing intra-residue distances and distances with a variance below the 95 % quantile in the contact matrix. By filtering based on the variance of the pairwise contact matrix we exclude long range distances and constant contacts from analysis. Consecutively the weighted arithmetic mean normalized mutual information matrix between all macroscopic features and atom-atom distances were computed with:

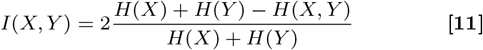

where *H*(·) is the information entropy of a feature and *H*(·, ·) is the joint information entropy of two features. Probability distributions to calculate the different entropies were estimated from min max scaled input features as re-weighted histograms. To ensure robustness in consecutive distance feature analysis the mutual information matrix was computed independently for each independent simulation replica and the considered features reduced to those passing following three criteria: (1) The distances average mutual information across simulation replicas with the funnel projection is above the 25 % quantile, (2) the distances relative standard deviation (*σ/µ*) of its mutual information across simulation replicas with the funnel projection is below the 25 % quantile, and (3) within all distances of a residue pair selected after the previous two filters select the distance maximising the mutual information with the macroscopic features. The reduced mutual information matrix of distances were clustered with MoSAIC using “modularity” (95). From the identified clusters only those with more than six member distances were selected as relevant motions correlated with the binding process in downstream analysis.

To assess how the selected correlated distances behave in relation to the binding event we cluster all simulation data along the binding process on the funnel projection with KMeans into a bound, transition, and unbound state. Re-weighted wasserstein distances were then computed between the unbound and bound state clusters. These distances were signed depending on a decrease or increase of the re-weighted distance value of highest probability between the unbound and bound state, respectively.

## Data, Materials, and Software Availability

Simulation input files (tpr), trajectories and colvar files are deposited in zenodo upon publication. Code used to generate plumed files, run simulations, and run analysis is available on https://gitlab.com/HankewieDanke/bh3_ligand_binding upon publication.

## ACKNOWLEDGMENTS

The authors would like to thank Dr. Nicola Piasentin, and Dr. Valerio Rizzi for discussion, proofreading and feedback on the manuscript.

## Footnotes

∗ Bcl-xL/BidBH3: 0.75; Bax/BidBH3: 0.5; Bax/*α*9: 0.4

