## Supplementary Figures and Tables for "How specific structural differences of Bcl2 proteins modulate the interaction with BH3 domains and apoptotic function"

**This PDF file includes:**

Supporting text
Figs. S1 to S17
Tables S1 to S7
SI References

### Supporting Information Text

### Re-folding of the Bcl-xL binding site

Initial simulations of Bcl-xL/BidBcl-2 homology (BH)3 peptide binding properly recovered a stronger $\Delta G_{bind}$ than Bax/BidBH3 peptide binding simulations, but the range of estimated $\Delta G_{bind}$ was large. One set presents a stronger $\Delta G_{bind}$ ($-21.96 \pm 0.43\, kcal/mol$, light blue) and the other a weaker $\Delta G_{bind}$ ($-13.48 \pm 0.65\, kcal/mol$, dark blue) (see Fig. S1 A, B). The former overestimates the experimental value, while the latter's estimate is close to the experimental values for Bcl-xL/tBid (see Tab. 1). Still, this weaker value also overestimates the affinity compared to experimental values of Bcl-xL/BidBH3 peptide affinity (see Tab. 1). This difference could be resulting from spontaneous gain of fold in the $\alpha3$ helix observed in the lower estimating simulation set, but not in the higher estimating simulation set (see Fig. S1).

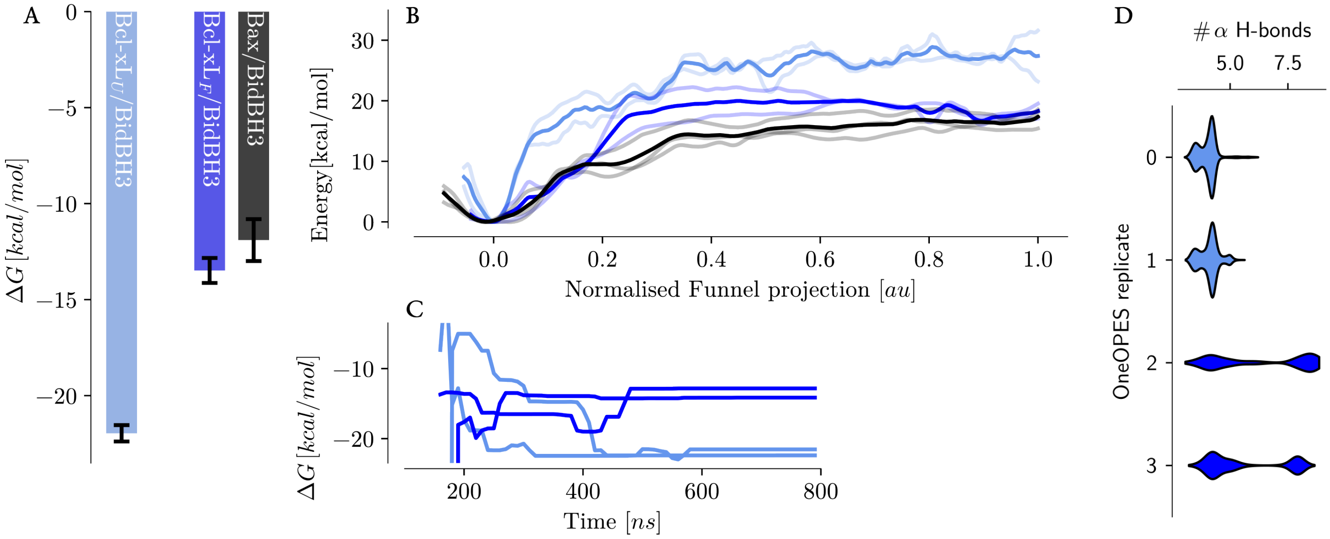


**Fig. S1.** Initial independent replicates (four) of enhanced sampling of BidBH3 peptide binding to Bcl-xL gave two different populations of $\Delta G_{bind}$ estimation. (A) Barplot of average $\Delta G_{bind}$ for the separate estimate populations, one strong estimation and one weak estimation. (B) Free energy surfaces along the funnel projection comparing the different estimate populations with the Bax/BidBH3 peptide system. (C) $\Delta G_{bind}$ estimates over time for the independent initial replicates. (D) Un-reweighted violinplots of the number of $\alpha-$helical hydrogen bonds in the $\alpha3$ helix of Bcl-xL in the independent OneOpes replicates generated for the Bcl-xL/BidBH3 peptide system. Note that replicate 0 and 1 do not show a subpopulation of high counts, while replicate 2 and 3 do. Replicate 0 and 1 have the stronger $\Delta G_{bind}$ estimates, while 2 and 3 have the weaker $\Delta G_{bind}$ estimates. Counts were calculated based on the distance between donor and acceptor atom transformed with a rational switching function (see Eq. 2, $N = 4, M = 14, R_0 = 5$ Å) and hydrogen bonds only considered, if donor and acceptor were four residues apart.

Binding simulations of BidBH3 peptide to Bcl-xL were thus re-run with an adjusted protocol taking the folding event within the binding site into account (see Tab. S5). This adjusted protocol biases the $\alpha$-helicity between residues R100 and P116 (see Eq. 5), through the $\alpha$-helical backbone hydrogen bonds present in the fully folded state but not present in the reference bound state (see Tab. S4). While still sampling the binding event within the correct ranking (see Fig. 2), this further allowed sampling of the fold gain in $\alpha3$ of Bcl-xL. Notably this lead to independent free energy surface (FES)'s with good agreement between them, where the fully folded state is a local minima and the predominantly unfolded state of the holo Bcl-xL/BidBH3 peptide is the global minima (see Fig. S2A, Fig. S7). Comparing the $\Delta G_{bind}$ estimate under consideration of the fold gain (see Tab. 1) with the estimates without any fold gain is in agreement with the estimated $\Delta G^{\alpha3}_{fold}$ ($\Delta G^{no-fold}_{bind} \sim \Delta G^{fold}_{bind} + \Delta G^{\alpha3}_{fold}$).

Comparing the fully folded and holo state of the $\alpha3$ helix in membrane vicinity with the solvent apo structure of Bcl-xL (PDB: 1LXL), we observe different fold states in all three (see Fig. S2B). The N-terminal side of $\alpha3$ contains a cluster of positive charge formed by arginines 100 to 103. In the solvent apo structure these residues are part of a loop pointing towards solvent, while the rest of $\alpha3$ remains helical and closes the binding site. In the membrane holo state these arginines become part of $\alpha2$ of Bcl-xL while the rest of $\alpha3$ is unfolded except for one turn, and in the fully-folded state instead become part of $\alpha3$. These Arginines show a high membrane contact within binding simulations (see Fig. S2C), and tend to be reduced when the $\alpha3$ helix is fully folded (see Fig. S3). Improved membrane contact of this Arginine cluster upon Bcl-xL$\alpha3$ re-folding, could be implicative of ligand helix binding improving the membrane association of the Bcl-xL globular domain.

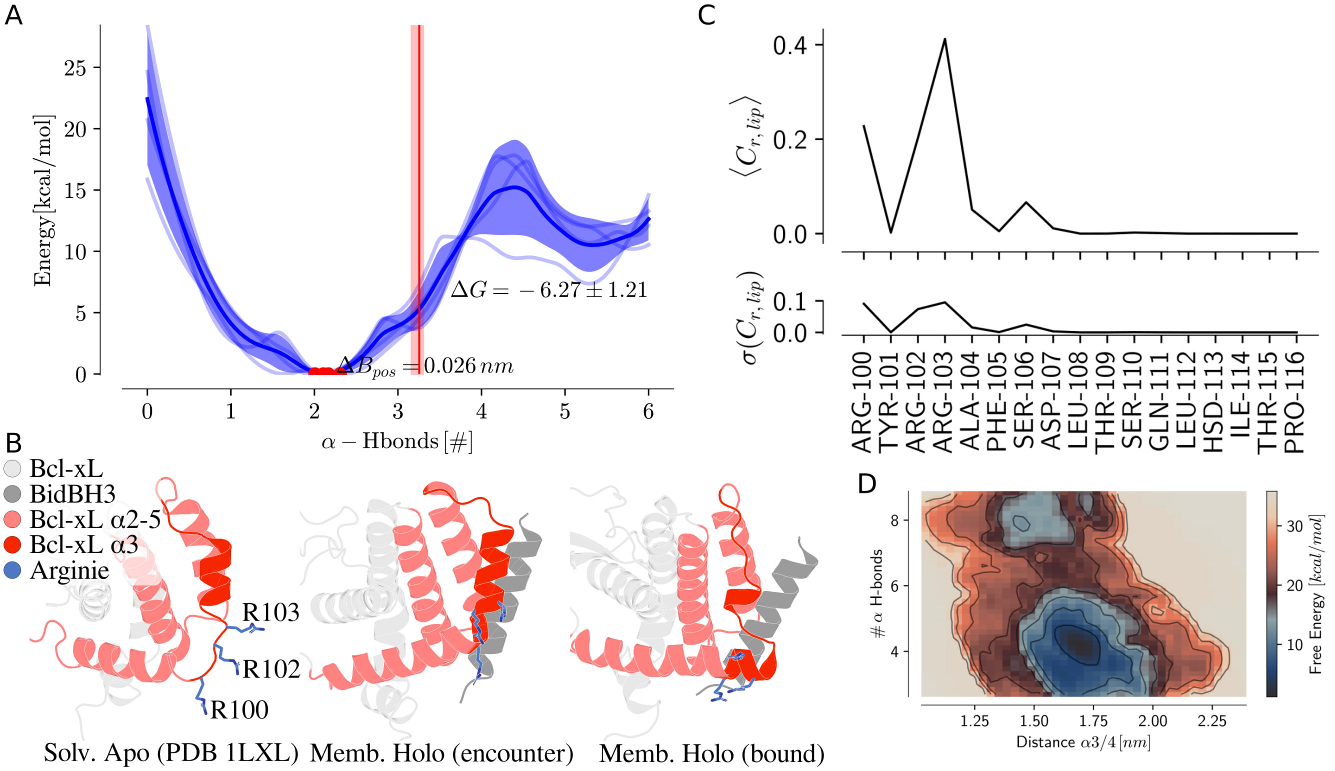


**Fig. S2.** $\alpha 3$ of Bcl-xL can take predominantly unfolded and fully folded states. (A) Free energy surface of fold-change within Bcl-xL's $\alpha 3$ helix along the Collective variable (CV) used as auxiliary variable. Individual independent replicates shown as transparent lines, average shown as line and standard deviation shown as transparent area. Global minima of the individual replicates shown as red dots and positioning standard deviation indicated as $\Delta B_{pos}$. Fold changing $\Delta G$ to the fully folded state indicated as text and state seperatrix indicated as red vertical line and shade. Seperatrices were automatically identified for independent replicates (see FES calculation and reweighting). Note that this count excludes hydrogen bonds present in the bound "un"-folded state. (B) Different fold states of Bcl-xL's $\alpha 3$ helix in different structural contexts. Bcl-xL shown as cartoon with membrane anchoring Arginines shown as sticks. The truncated ($\Delta\alpha 9$) solvent apo structure is PDB entry 1LXL (1). (C) Average non-reweighted membrane contact of residues in $\alpha 3$ with the closest lipid (top) and their variance (bottom) across all independent binding simulation replicates. Contacts were calculated with a tanh switching function ($C = -0.5\tanh(10d_{r,l} - 5) - 1$) and between the center of mass of protein residues and lipid phosphates. (D) Two dimensional FES of the binding site accessibility (distance between $\alpha 3$&4), and $\alpha 3$ helicity. This shows a slight negative correlation, where the binding site distance decreases upon increase of helicity. Here the number of $\alpha$ hydrogen bonds were computed between all residues between 100 and 116.

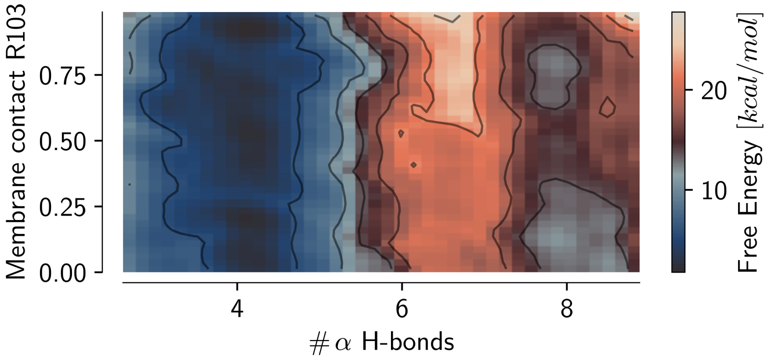


**Fig. S3.** Two dimensional FES of the Bcl-xL/BidBH3 peptide system in the membrane contact between Bcl-xL $R103$ and $\alpha 3$ helicity. FES calculated as average across the four independent simulation replica. Helicity is defined as the number of all $\alpha$-helical hydrogen bonds (see Eq. 5) in $\alpha 3$. Membrane contact is the contact between $R103$ and the closest lipid defined as in Fig. S2.

### Figures and Tables

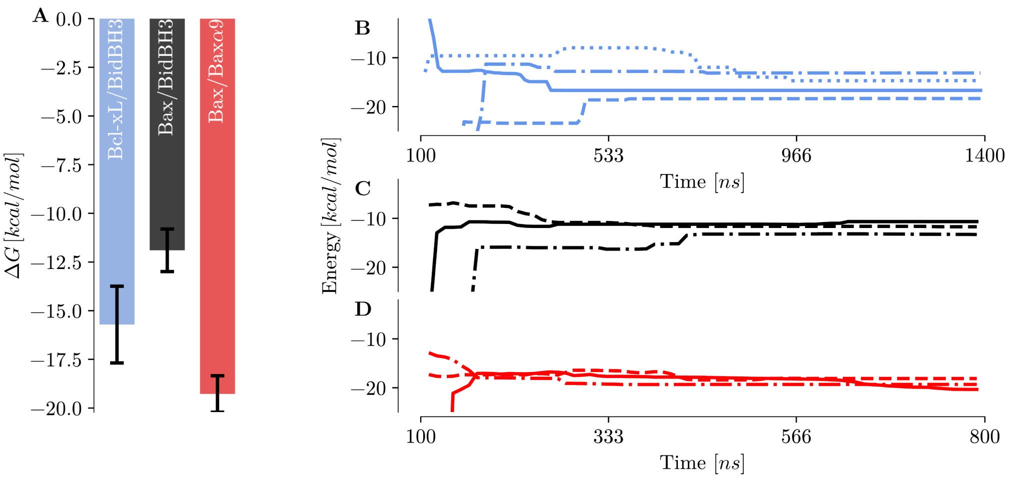


**Fig. S4.** Funnel FES's estimated $\Delta G_{bind}$ values. (A) Barplot of estimated final $\Delta G_{bind}$ values over all considered simulation timesteps. (B, C, D) $\Delta G_{bind}$ between bound and unbound state over time for the different simulation systems. Separate line types for independent replicates.

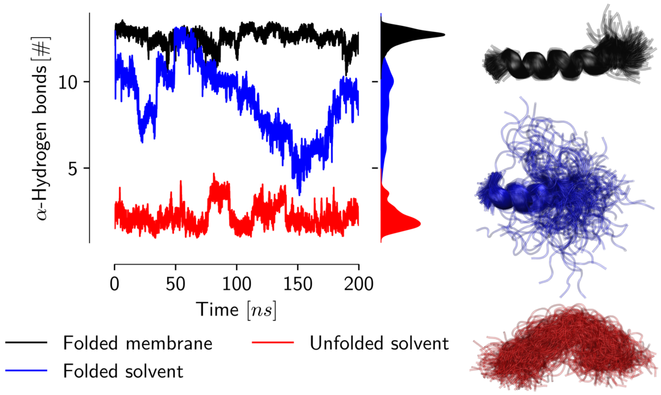


**Fig. S5.** The BidBH3 peptide helix is unstable in its unbound solvent state. BidBH3 peptide helices unfold when simulated in plain MD within pure solvent. Helices were simulated (equilibration and production) in TIP3P water either starting from a folded state (blue), or starting from an unfolded state (red). When a membrane is present the folded state is stable (over $200\,ns$, black). The unfolded state was obtained by breaking all backbone hydrogen bonds of the helix via adiabatic bias molecular dynamics (MD).

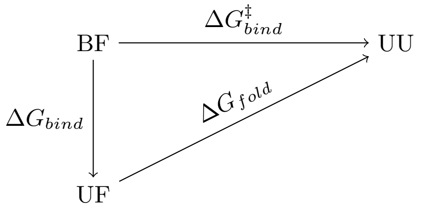


$$\Delta G^{\ddagger}_{bind} = \Delta G_{bind} + \Delta G_{fold}$$

**Fig. S6.** Correction term of the binding estimate derives from the separated binding and folding processes. This goes through an intermediate state from bound folded (BF) to unbound unfolded (UU) through unbound folded (UF).

 Anton Hanke, Christina Elsner, Simone Aureli, Enrica Bordignon, Francesco Luigi Gervasio

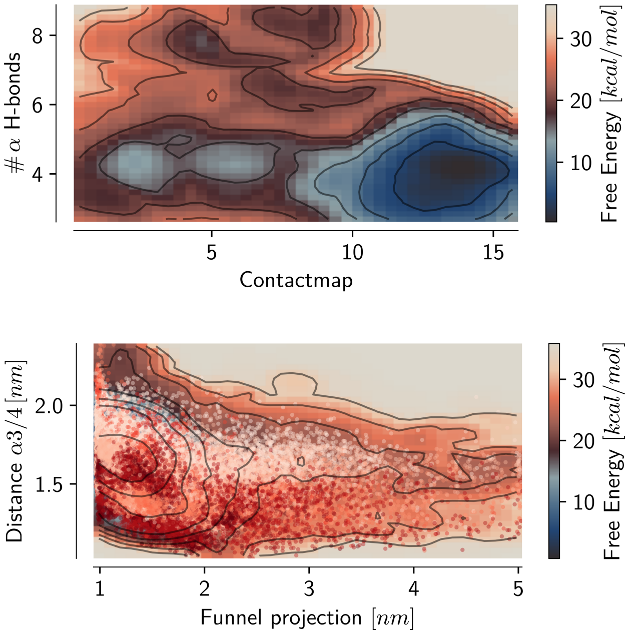


**Fig. S7.** Two dimensional FES of the Bcl-xL/BidBH3 peptide system in the contactmap and $\alpha3$ helicity (top) and in the funnel projection and distance between Bcl-xL $\alpha3$ and $\alpha4$ with data-points shown as scatter colored by $\alpha3$ helicity (bottom). While the funnel projection suggests, that the $\alpha3$ helix of Bcl-xL can gain fold within the bound state, the contactmap shows that the fully bound state becomes unreachable upon fold gain. FES calculated as average across the four independent simulation replica. Helicity is defined as the number of all $\alpha$-helical hydrogen bonds (see Eq. 5) in $\alpha3$.

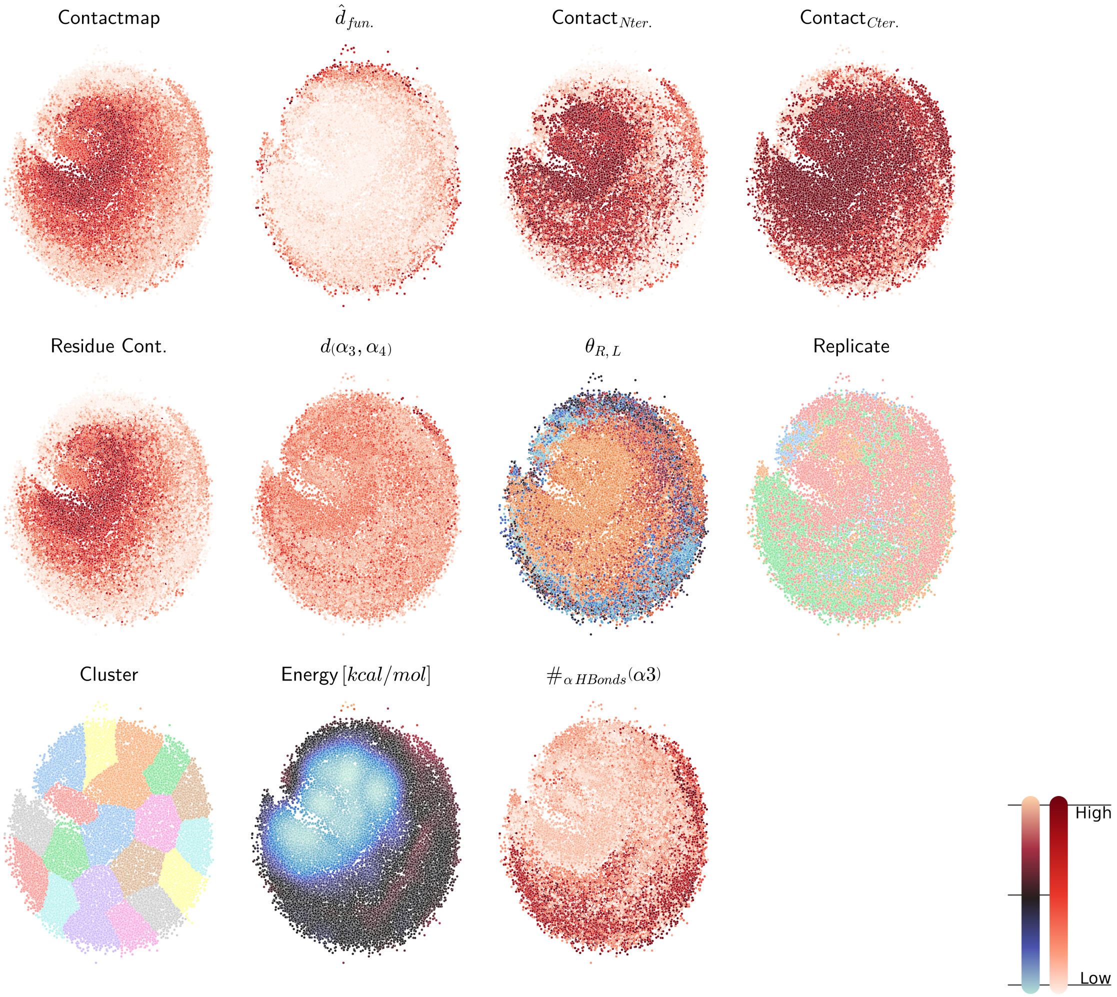


**Fig. S8.** ELViM dimensionality reduction of the Bcl-xL/BidBH3 peptide simulations. Conformational ensemble of all independent replicates plotted within the ELViM dimensionality reduction, coloured by different simulation features, energy, and extracted clusters. Colour schemes: (bright to dark red) low to high value, (light blue to light red/orange) low to high value, (discrete colours) coloured by index. PyMOL sessions of cluster representatives are available online.

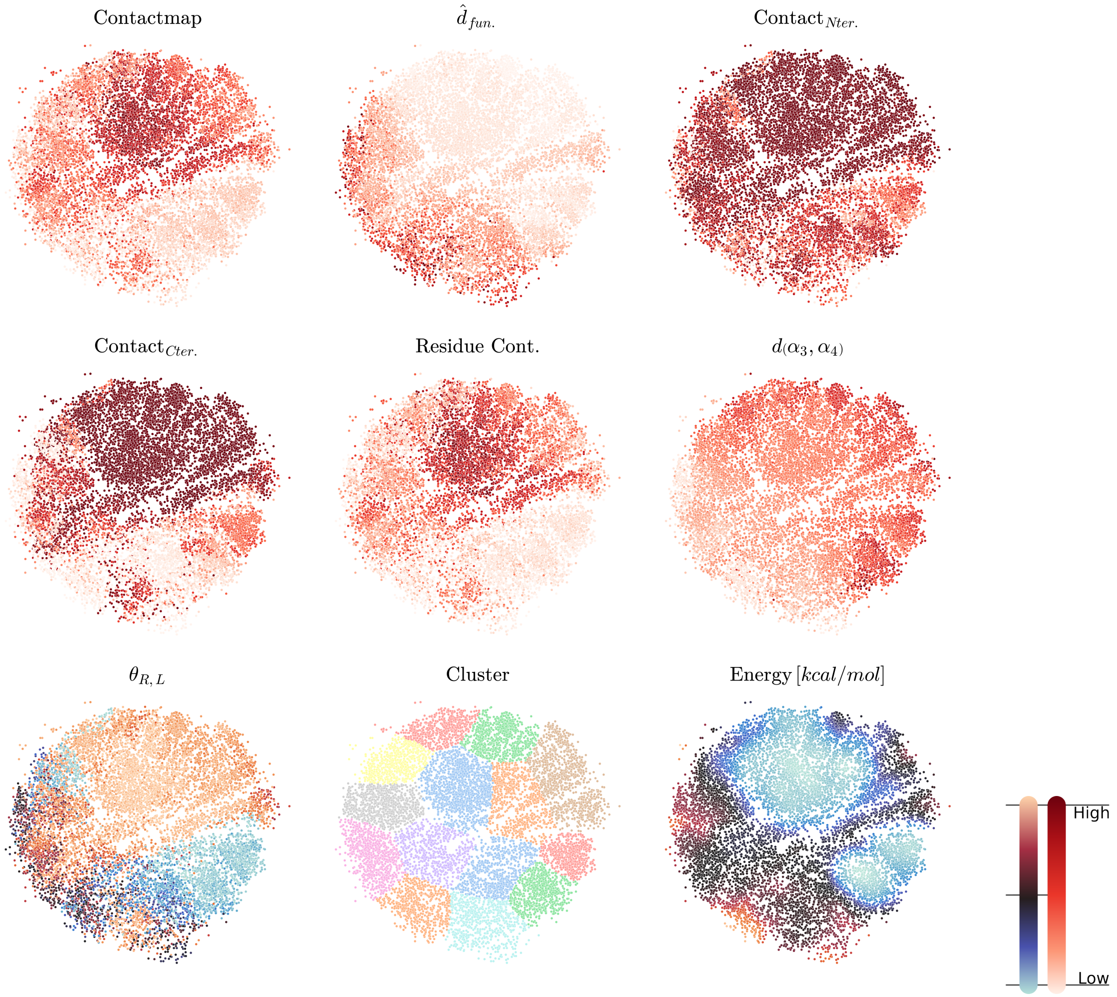


**Fig. S9.** ELViM dimensionality reduction of the Bax/BidBH3 peptide simulations. Conformational ensemble of all independent replicates plotted within the ELViM dimensionality reduction, coloured by different simulation features, energy, and extracted clusters. Colour schemes: (bright to dark red) low to high value, (light blue to light red/orange) low to high value, (discrete colours) coloured by index. PyMOL sessions of cluster representatives are available online.

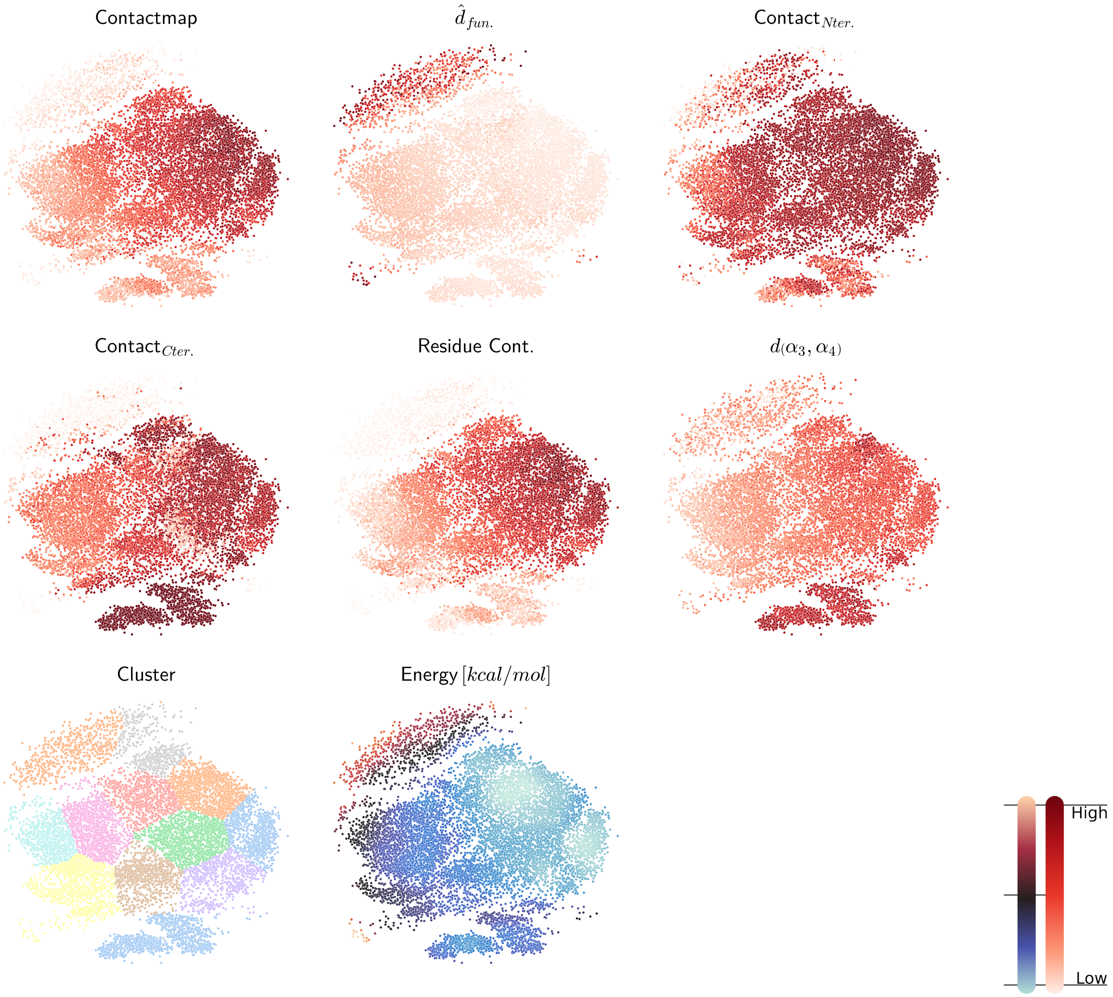


**Fig. S10.** ELViM dimensionality reduction of the Bax/$\alpha 9$ simulations. Conformational ensemble of all independent replicates plotted within the ELViM dimensionality reduction, coloured by different simulation features, energy, and extracted clusters. Colour schemes: (bright to dark red) low to high value, (light blue to light red/orange) low to high value, (discrete colours) coloured by index. PyMOL sessions of cluster representatives are available online.

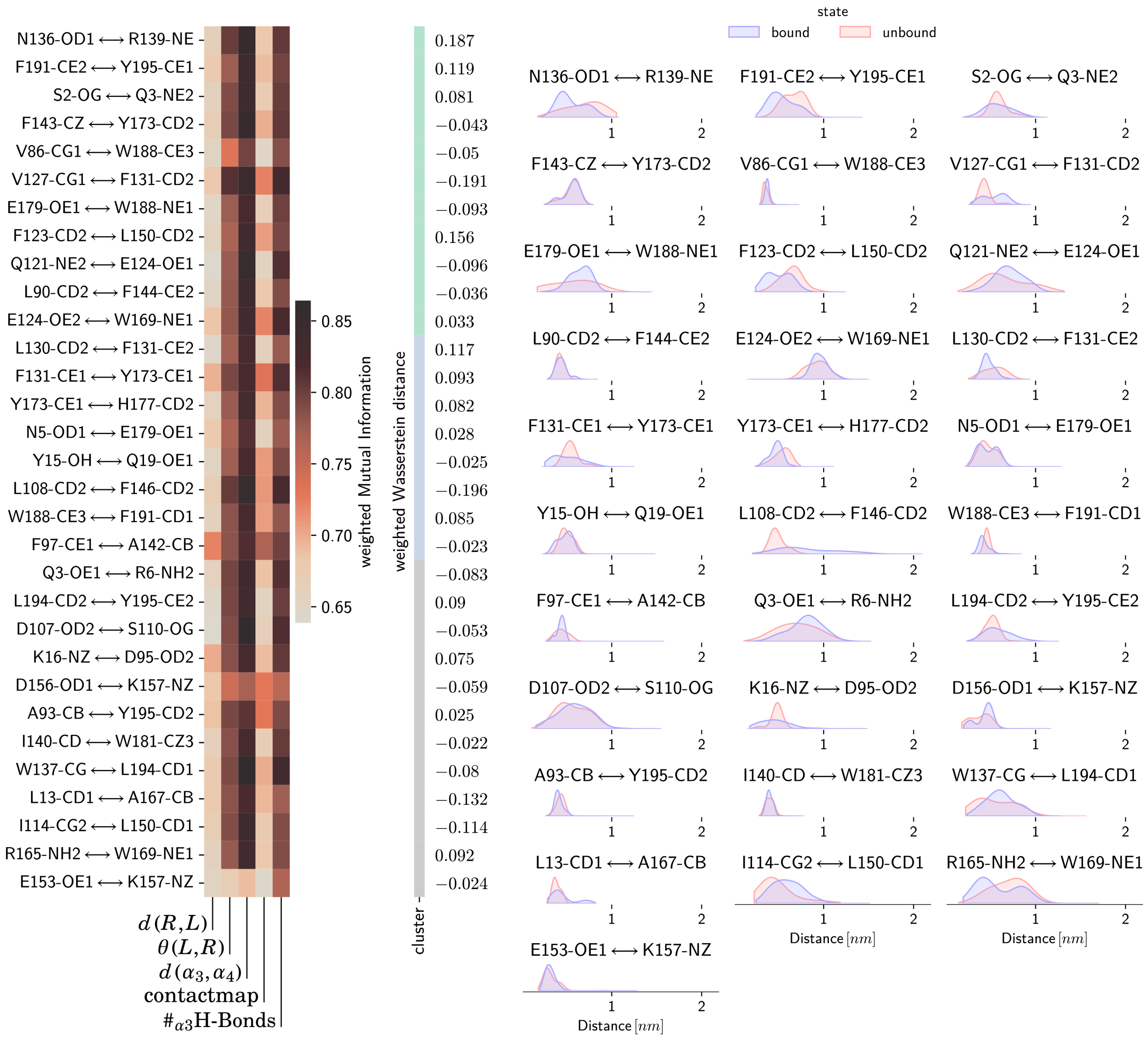


**Fig. S11.** Mutual Information analysis of residue contacts with macroscopic features in the Bcl-xL/BidBH3 peptide simulations. Heatmap shows the re-weighted average normalized mutual information between atom-atom distances and macroscopic features such as the ligand/receptor contactmap and binding site distance ($d\ (\alpha 3,\ \alpha 4)$). Cluster column of the heatmap indicates mutual information clusters between atom-atom distances and is labeled by the signed re-weighted wasserstein distance between bound and unbound state of the ligand ($d_{ws} > 0$ distance increase from bound to unbound). Distribution plots show the re-weighted distance distributions between unbound and bound states identified based on the ligand/receptor contactmap. PyMOL sessions visualising this analysis are available online.

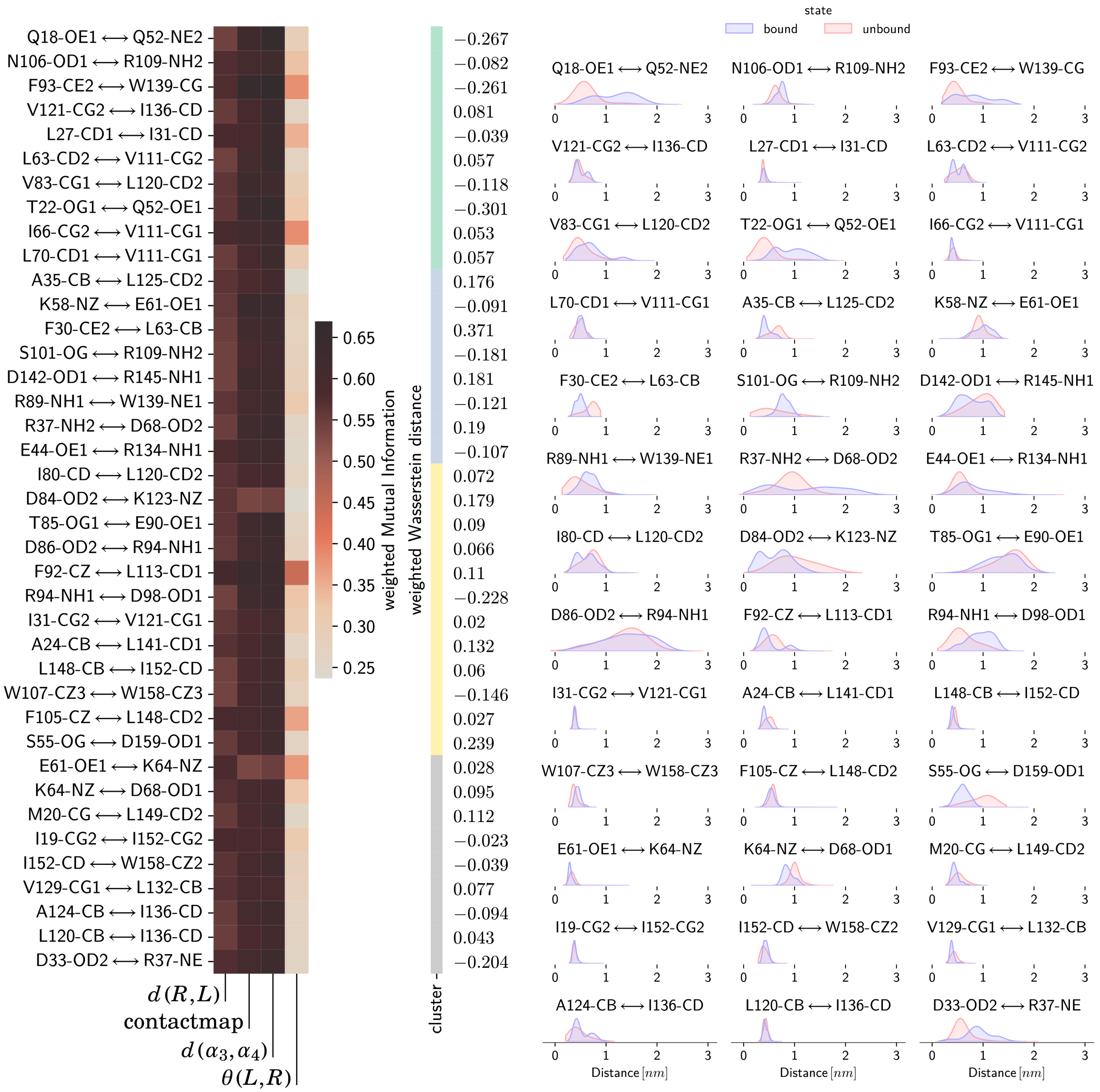


**Fig. S12.** Mutual Information analysis of residue contacts with macroscopic features in the Bax/BidBH3 peptide simulations. Heatmap shows the re-weighted average normalized mutual information between atom-atom distances and macroscopic features such as the ligand/receptor contactmap and binding site distance ($d\,(\alpha 3, \alpha 4)$). Cluster column of the heatmap indicates mutual information clusters between atom-atom distances and is labeled by the signed re-weighted wasserstein distance between bound and unbound state of the ligand ($d_{ws} > 0$ distance increase from unbound to unbound). Distribution plots show the re-weighted distance distributions between unbound and bound states identified based on the ligand/receptor contactmap. PyMOL sessions visualising this analysis are available online.

 

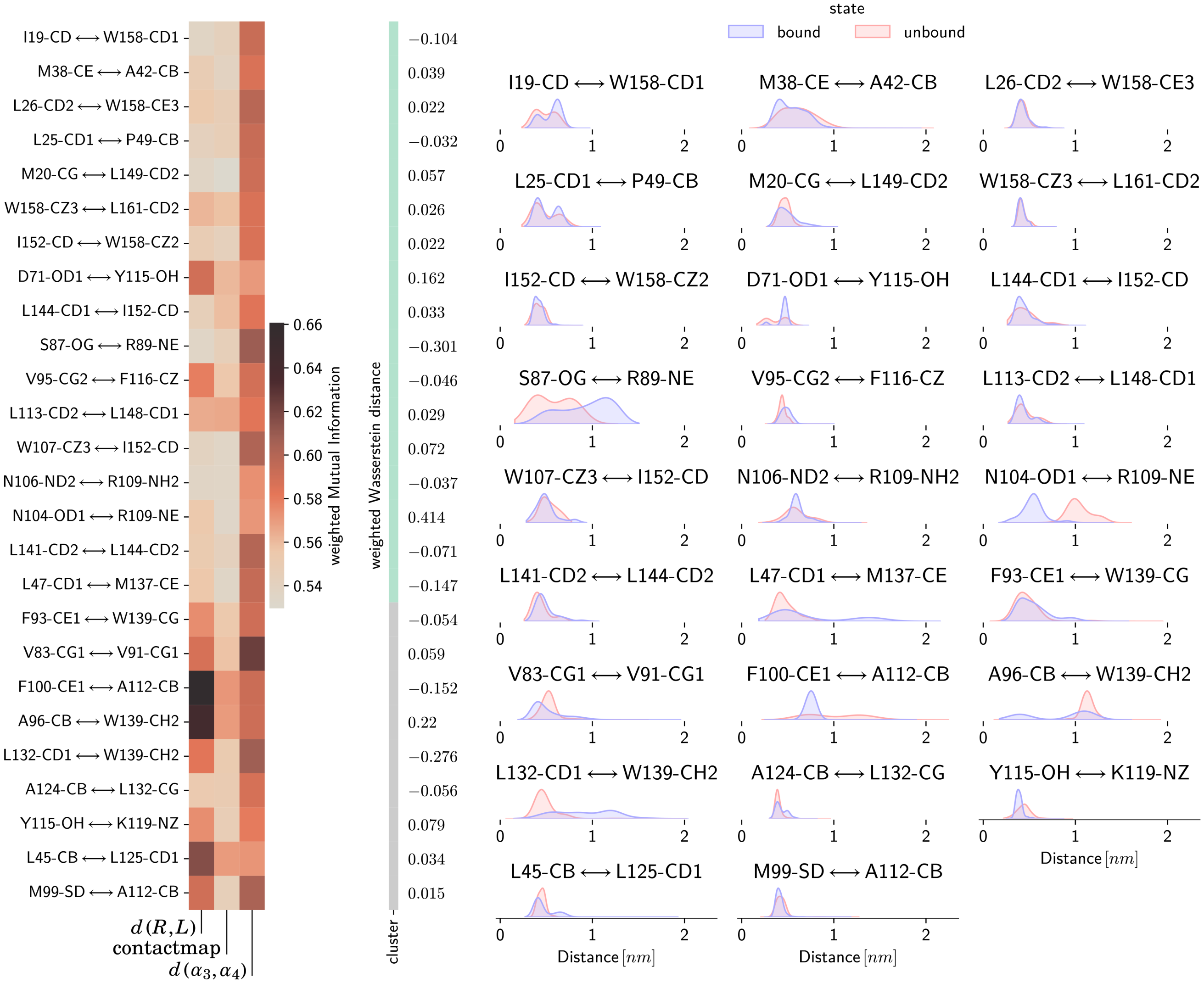


**Fig. S13.** Mutual Information analysis of residue contacts with macroscopic features in the Bax/$\alpha9$ simulations. Heatmap shows the re-weighted average normalized mutual information between atom-atom distances and macroscopic features such as the ligand/receptor contactmap and binding site distance ($d\,(\alpha3, \alpha4)$). Cluster column of the heatmap indicates mutual information clusters between atom-atom distances and is labeled by the signed re-weighted wasserstein distance between bound and unbound state of the ligand ($d_{ws} > 0$ distance increase from bound to unbound). Distribution plots show the re-weighted distance distributions between unbound and bound states identified based on the ligand/receptor contactmap. PyMOL sessions visualising this analysis are available online.

**Table S1. Charges of protein models. Note that both PDB2PQR and CHARMM-GUI use propka for pKa assignment and protonation.**

| System | Charge $[e]$ | Tool |
|---|---:|---|
| 1f16(2) | -3 | |
| Apo Bax | -3 | PDB2PQR(3, 4) |
| Apo BclxL | -11 | |
| Holo Bax | -5 | |
| Holo Bcl-xL | -13 | CHARMM-GUI(5) |
| Apo BidBH3 peptide | -2 | |

**Table S2. System compositions. Systems were set up with GROMACS in solvent and CHARMM-GUI when a membrane is present.**

| System | Tool | Lipids (CL/POPC) | Ions (Na/Cl) | Waters | Atoms |
|---|---|---|---|---|---|
| 1f16(2) | gromacs | — | 89/86 | 30176 | 93680 |
| BidBH3 peptide | | — | 25/23 | 8221 | 25033 |
| Holo Bax | CHARMM-GUI | 42/164 (20/80 %) | 147/58 | 21668 | 100565 |
| Holo Bcl-xL | | 60/240 (20/80 %) | 253/120 | 43607 | 181659 |
| BidBH3 peptide | | 15/60 (20/80 %) | 44/12 | 4723 | 26187 |

**Table S3. Simulation protocol and restraint used throughout. For detailed description on other simulation parameters see .**

| System | integrator | nsteps | emtol | constraints | constraint algorithm | replicates |
|---|---|---|---|---|---|---|
| Membrane | steep | 5000 | 1000.0 | h-bonds | LINCS | 1 |
| Solvent | steep | 5000 | 1000.0 | h-bonds | LINCS | 1 |

| Membrane | Step | Ensemble | Duration $[ns]$ | Restraints $[kJ/mol]$ Sidechain | Backbone | Lipid Z | Lipid dihed | Continue | Repl |
|---|---|---|---|---|---|---|---|---|---|
| | 0 | NVT | 0.125 (1fs) | 4000 | 2000 | 1000 | 1000 | No | 1 |
| | 1 | NVT | 0.125 (1fs) | 2000 | 1000 | 400 | 400 | No | 1 |
| Equil. | 2 | NPT | 0.125 (1fs) | 1000 | 500 | 400 | 200 | No | 1 |
| | 3 | NPT | 0.5 (2fs) | 500 | 200 | 200 | 200 | No | 1 |
| | 4 | NPT | 0.5 (2fs) | 500 | 200 | 200 | 200 | No | 1 |
| Prod. | Bcl-xL (4.1) | NPT | 100.0 (2fs) | 0 | 0 | 0 | 0 | No | 1 |
| | Bax (4.1) | NPT | 50.0 (2fs) | 0 | 0 | 0 | 0 | No | 1 |

| Solvent Bax | Step | Ensemble | Duration $[ns]$ | Restraints $[kJ/mol]$ Sidechain | Backbone | Continue | Repl |
|---|---|---|---|---|---|---|---|
| | 0 | NVT | 0.125 (1fs) | 1000 | 200 | No | 1 |
| | 1 | NVT | 0.125 (1fs) | 1000 | 200 | No | 1 |
| Equil. | 2 | NPT | 0.125 (1fs) | 1000 | 200 | No | 1 |
| | 3 | NPT | 0.5 (2fs) | 0 | 200 | No | 1 |
| Prod. | 3.1 | NPT | 100.0 (2fs) | 0 | 0 | No | 1 |

| BidBH3 (+mem.) | Step | Ensemble | Duration $[ns]$ | Restraints $[kJ/mol]$ Backbone H-Bonds | Continue | Repl |
|---|---|---|---|---|---|---|
| Equil. | 0 | NVT | 10 (2fs) | 1000 | No | 1 |
| | 1 | NPT | 10 (2fs) | 1000 | No | 1 |
| ABMD unfolding | 2 | NPT | COMMITOR (2fs) | $\kappa_{ABMD} = 2$ | No | 1 |
| Prod. unfolded | 3.0 | NPT | $\sim 200$ (2fs) | — | No | 1 |
| Prod. folded | 3.1 | NPT | $\sim 200$ (2fs) | — | No | 1 |
| Prod. mem. folded | 3.2 | NPT | $\sim 200$ (2fs) | — | No | 1 |

**Table S4. Auxiliary CVs biased during simulations.**

| CV | Description | Systems |
|---|---|---|
| pp.proj | Distance along the funnel projection. Biased to increase sampling in bulk solvent. | ● (blue) |
| Water Coord. | Coordination of the binding site center of mass (CoM) with water. Help with displacement of ligand from the binding site by actively increasing the water density in the binding site. | ● (red) ● (black) ● (blue) |
| Ligand/receptor angle | Angle between the vector along the ligand helix and along the binding site (biased within bulk solvent). | ● (red) ● (black) ● (blue) |
| $\#\alpha_3$-HBonds | Sum of backbone hydrogen bonds forming the $\alpha$-helix 3 of Bcl-xL that are not present in the fully bound state, but are in the folded state of the helix. The sum was calculated as in equation 5. | ● (red) |

● (black) Bax/BidBH3, ● (red) Bcl-xL/BidBH3, ● (blue) Bax/$\alpha9$

**Table S5. OneOpes setup of OPES biases used within the binding simulations.**

| System | OPES Flavour | Biased CV(s) | PACE | BARRIER | SIGMA | Replicate 0 | 1 | 2 | 3 | 4 | 5 | 6 | 7 |
|---|---|---|---|---|---|---|---|---|---|---|---|---|---|
| Bax/BH3 | Explore | cv, pp.proj | 10000 | 60 | 0.1,0.2 | • | • | • | • | None | | | |
| | | Water | 10000 | 30 | 0.2 | | • | • | • | | | | |
| | | $\Theta_{R,L}$, pp.proj | 10000 | 30 | 0.1, 0.1 | | | • | • | | | | |
| Initial BclxL/BH3 | Explore | cv, pp.proj | 10000 | 85 | 0.1,0.2 | • | • | • | • | None | | | |
| | | Water | 10000 | 20 | 0.2 | | • | • | • | | | | |
| | | $\Theta_{R,L}$, pp.proj | 10000 | 30 | 0.1, 0.1 | | | • | • | | | | |
| Final BclxL/BH3 | Explore | cv, pp.proj | 10000 | 100 | 0.1,0.2 | • | • | • | • | None | | | |
| | | $\#\alpha_3$-HBonds | 10000 | 40 | 0.1 | | • | • | • | | | | |
| | | $\Theta_{R,L}$, pp.proj | 10000 | 30 | 0.1, 0.1 | | | • | • | | | | |
| | | Water | 10000 | 20 | 0.2 | | | | • | | | | |
| Bax/$\alpha 9$ | Explore | cv | 10000 | 90 | 0.1 | • | • | • | • | None | | | |
| | | Water | 10000 | 10 | 0.2 | | • | • | • | | | | |
| | | pp.proj | 10000 | 10 | 0.1 | | • | • | • | | | | |
| | | $\Theta_{R,L}$ | 10000 | 3 | 0.1 | | | • | • | | | | |

**Table S6. OneOpes setup of OPES biases used within the BidBH3 helix folding.**

| System | OPES Flavour | Biased CV(s) | PACE | BARRIER | SIGMA | Replicate 0 | 1 | 2 | 3 | 4 | 5 | 6 | 7 |
|---|---|---|---|---|---|---|---|---|---|---|---|---|---|
| BidBH3 helix | Explore | diffHB, cmap | 10000 | 100 | 1.46,0.46 | ● | ● | ● | ● | ● | ● | ● | ● |
| | | C195 | 10000 | 30 | 0.32 | | ● | ● | ● | ● | ● | ● | ● |
| | | C125 | 10000 | 30 | 0.3 | | | ● | ● | ● | ● | ● | ● |
| | | C144 | 10000 | 30 | 0.24 | | | | ● | ● | ● | ● | ● |
| | | C115 | 10000 | 30 | 0.28 | | | | | ● | ● | ● | ● |
| | | O69 | 10000 | 30 | 0.31 | | | | | | ● | ● | ● |
| | | N113 | 10000 | 30 | 0.22 | | | | | | | ● | ● |
| | | O196 | 10000 | 30 | 0.43 | | | | | | | | ● |
| | Multithermal | Energy | 1000 | | | | 315.00 | 320.00 | 325.00 | 330.00 | 335.00 | 340.00 | 345.00 |

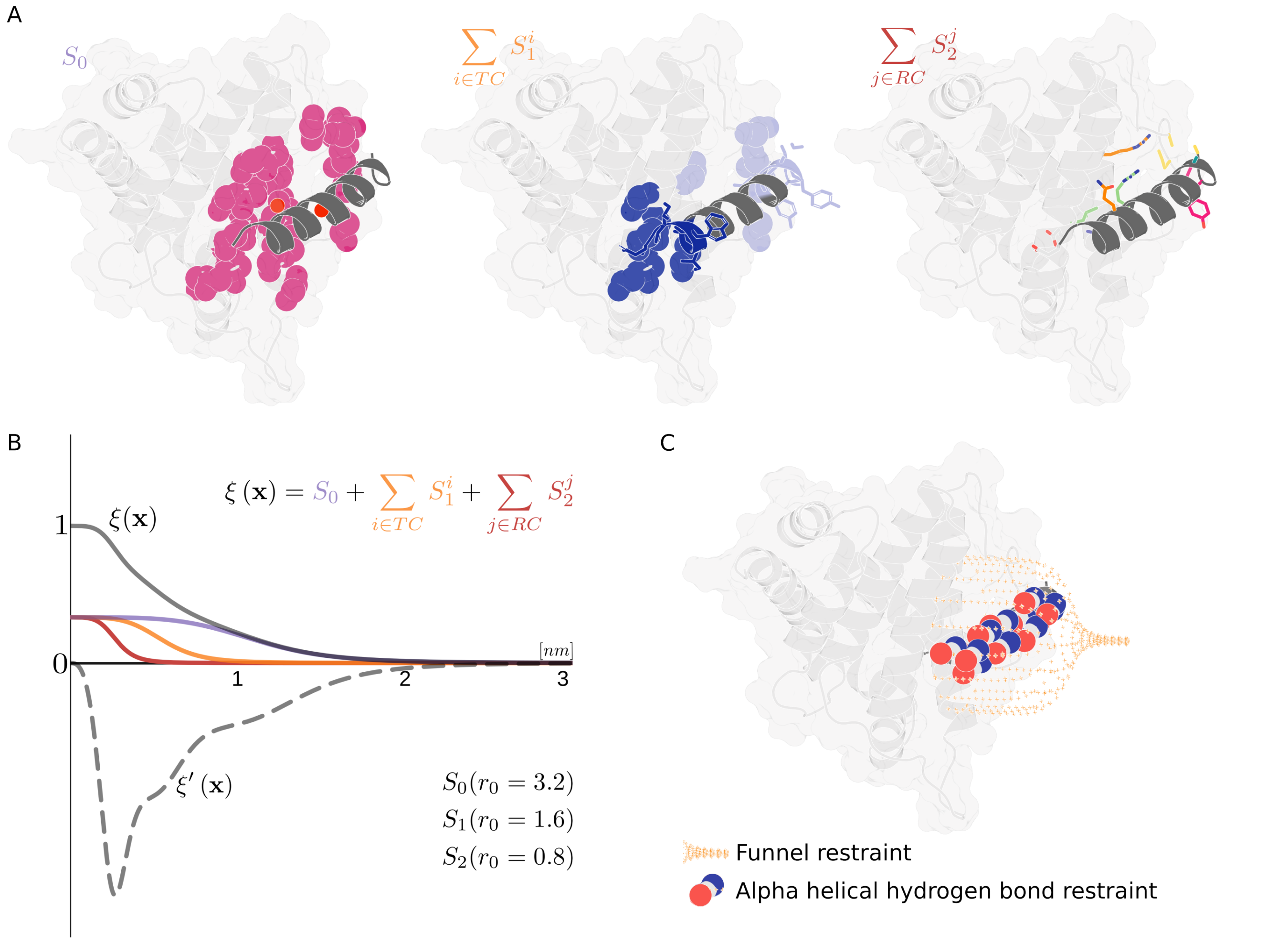


**Fig. S14.** Main Collective variable ($\xi$) biased and restraints applied. (A) Cartoon and surface representation of the multi level contact map. Left: long range interaction defined between center of mass (red spheres) of ligand and binding groove (coloured spheres). Center: intermediate range interactions defined between center of mass of ligand N- and C-terminus to their respective region of the binding groove (coloured spheres). Right: Residue-residue contacts defined by center of mass distances between side-chains of interacting residues. For each ligand residue in the interface, binding groove selection consists of their direct contacts (sticks). (B) Line plots on the values of separate switching functions associated with contacts, depending on their nature in (A). Sum of contacts ($\xi\ (x)$) shown in grey, if one contact of each type exists (TC: terminal contact, RC: residue contact). Derivative of the CV ($\xi'\ (x)$) shown as grey dashed line. (C) Main restraints applied during simulations, cartoon representation of ligand and receptor and spheres of atoms involved in backbone hydrogen bonds of the ligand. Funnel-restraint indicated with crosses.

**Table S7. Additional restraints needed.**

| Restraint | Description | $\kappa\ [kJ/mol]$ | Systems |
|---|---|---|---|
| Funnel to membrane plane | The Bcl-xL globular domain is weakly anchored to the membrane, as such many conformations can result in the funnel pointing towards the membrane. By restraining the Angle of the funnel axis to the z-axis to $90°$ the funnel continues pointing towards solvent. | 200 | ● (red) |
| Bcl-xL globular domain to membrane | The z-distance between the upper membrane leaflet and Bcl-xLs globular domain is CoM restrained to avoid biases in the system detaching the domain from the membrane. | 200 | ● (red) |
| Binding site helicity | The binding sites showed some tendency to unfold in $\alpha 3-4$ (canonical binding site) and were restrained via the backbone hydrogen bonds in helices $\alpha 3-4$ (see Eq. 5), which are present in the bound state. | 30 | ● (black) ● (red) ● (blue) |
| Funnel projection to multi level contact-map | To dissuade unfolding and sampling of non-physical conformations initial simulations were used to fit logarithmic functions: $\hat{d}_e(x) = \sum_{i=0}^{2} n_i \log(S(x))^{3-i} + c$. The deviation from this logarithmic polynomial was upper wall constrained towards $0.5 \geq \hat{d}(x) - \hat{d}_e(x)$ with differing $\kappa$ dependent on the simulation system. | 300/1200 | ● (black) ● (red) |

● (black) Bax/BidBH3, ● (red) Bcl-xL/BidBH3, ● (blue) Bax/$\alpha 9$

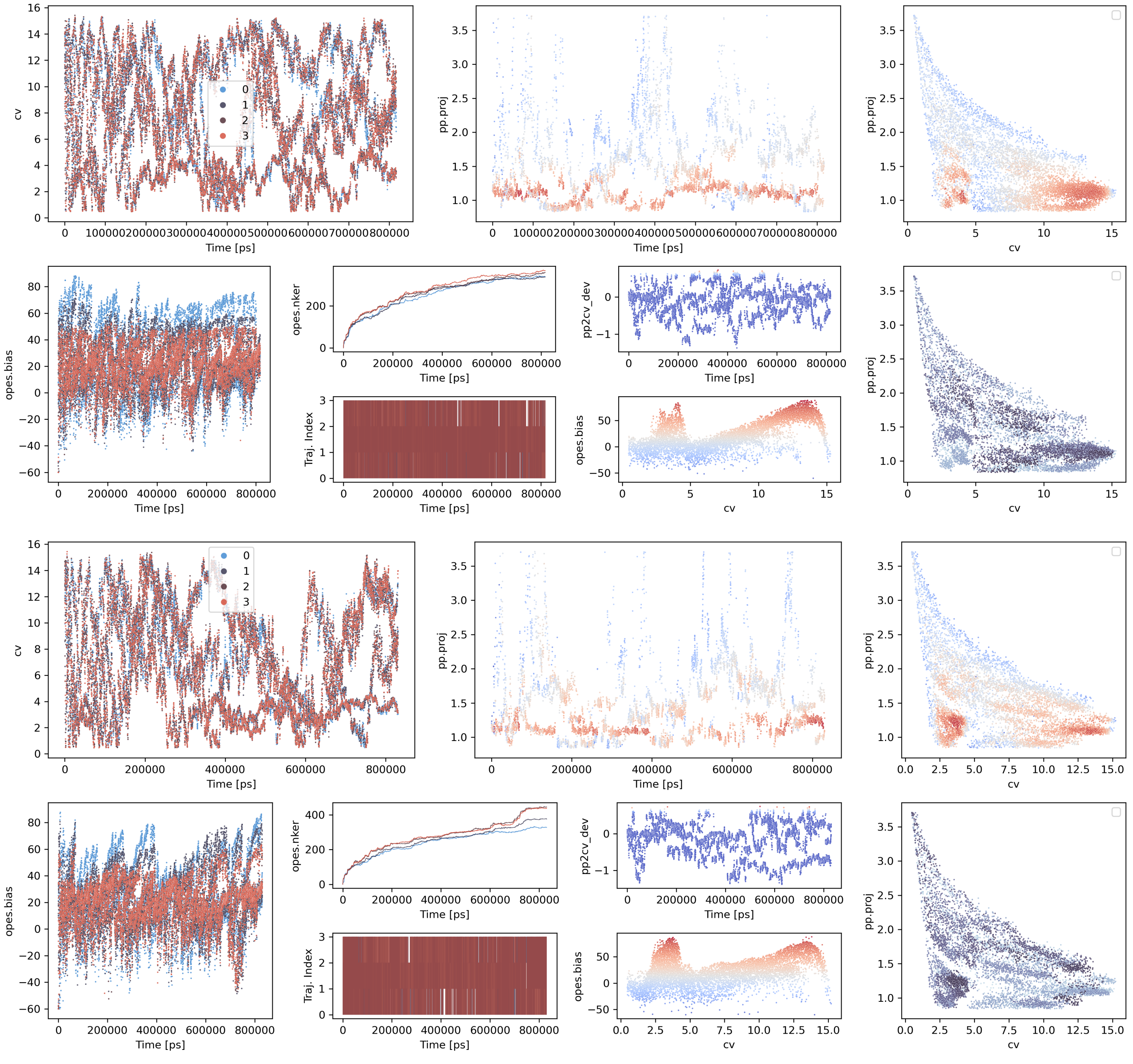


**Fig. S15. Simulation traces of three OneOPES simulations for the Bax/BidBH3 peptide system.** One set of plots per technical replicate, each set containing two rows. Top row, left to right: (1) contactmap value over time colored by replicate; (2) funnel projection over time for the base replicate colored by bias; (3) Contactmap against funnel projection colored by bias in the base replica. Bottom row, left to right: (4) Bias of the main opes call colored by replicate, (5, top) Number of kernels in the main CV opes colored by replicate, (5, bottom) trajectory index within exchange over time colored by replicate, (6, top) deviation from logarithmic fit between contactmap and funnel projection from test simulations, (6, bottom) Main bias over the contactmap colored by bias, (7) contactmap against funnel projection colored by time in the base replica.

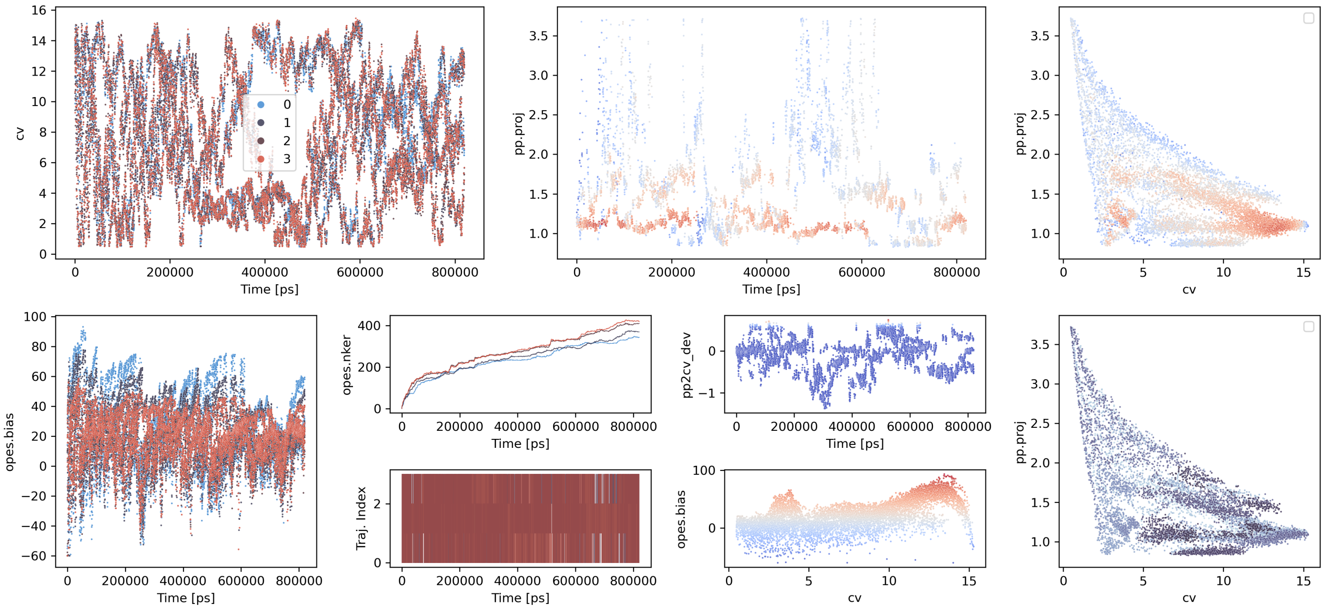


**Fig. S15. Simulation traces of three OneOPES simulations for the Bax/BidBH3 peptide system.** (continuation)

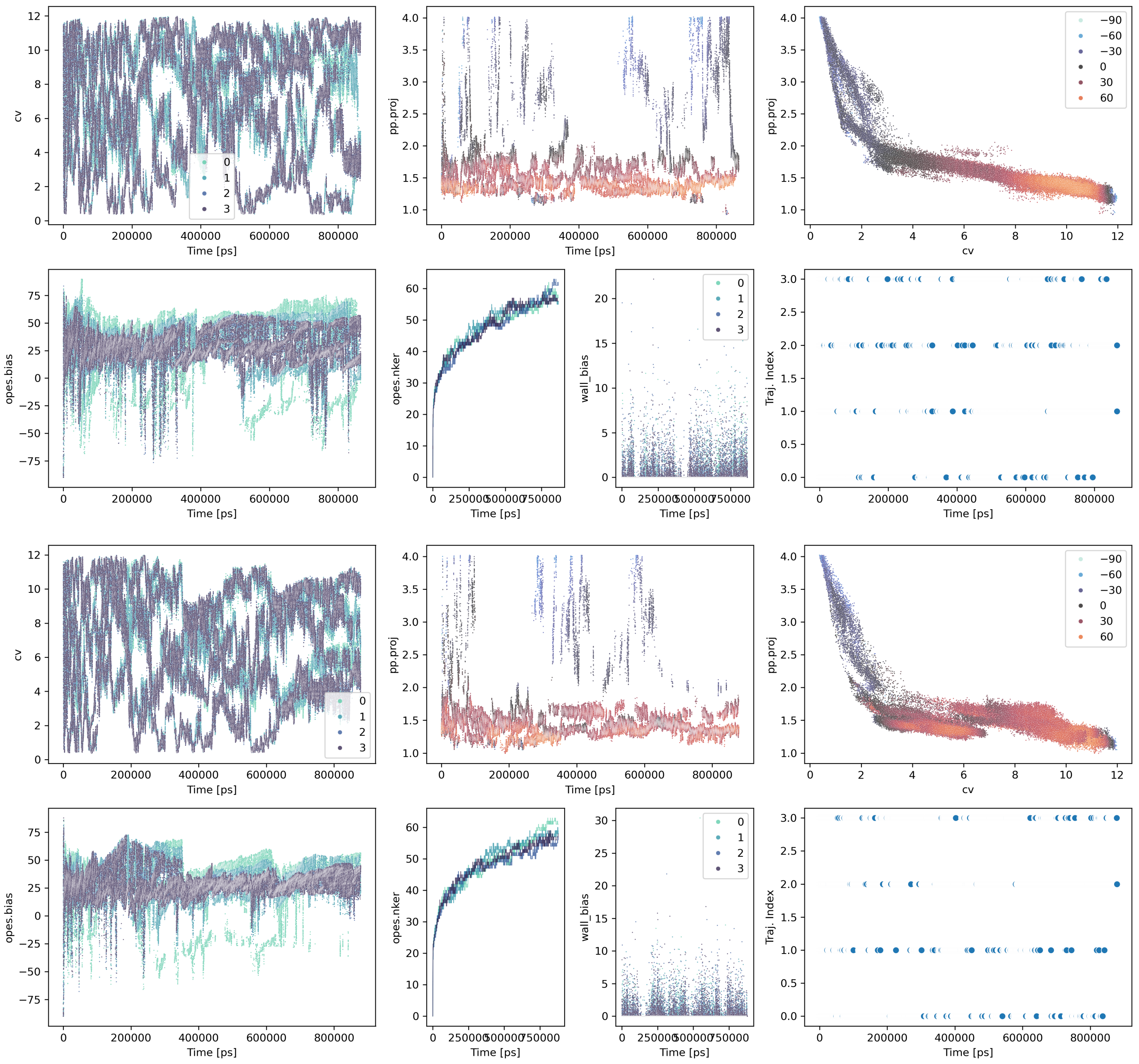


**Fig. S16. Simulation traces of three OneOPES simulations for the Bax/$\alpha 9$ system.** One set of plots per technical replicate, each set containing two rows. Top row, left to right: (1) contactmap value over time colored by replicate; (2) funnel projection over time for the base replicate colored by bias; (3) Contactmap against funnel projection colored by bias in the base replica. Bottom row, left to right: (4) Bias of the main opes call colored by replicate, (5) Number of kernels in the main CV opes colored by replicate, (6) bias applied by the funnel wall across all replicates, (7) Trajectory index over time of the base replicate.

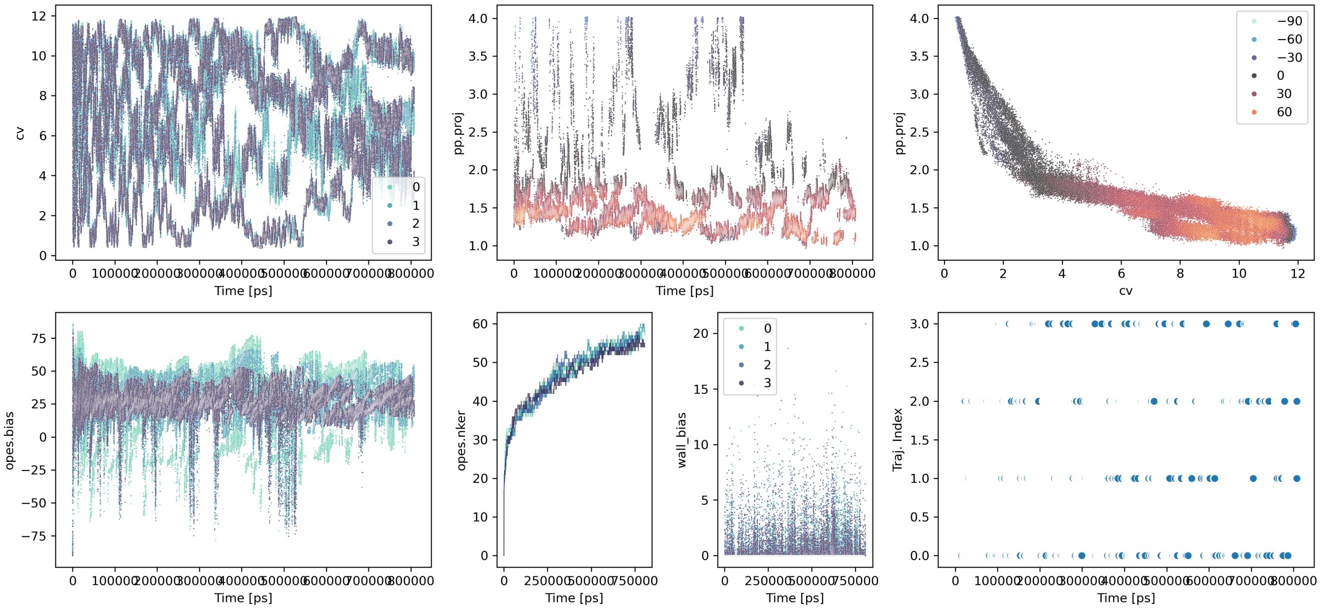


**Fig. S16. Simulation traces of three OneOPES simulations for the Bax/$\alpha 9$ system.** (continuation)

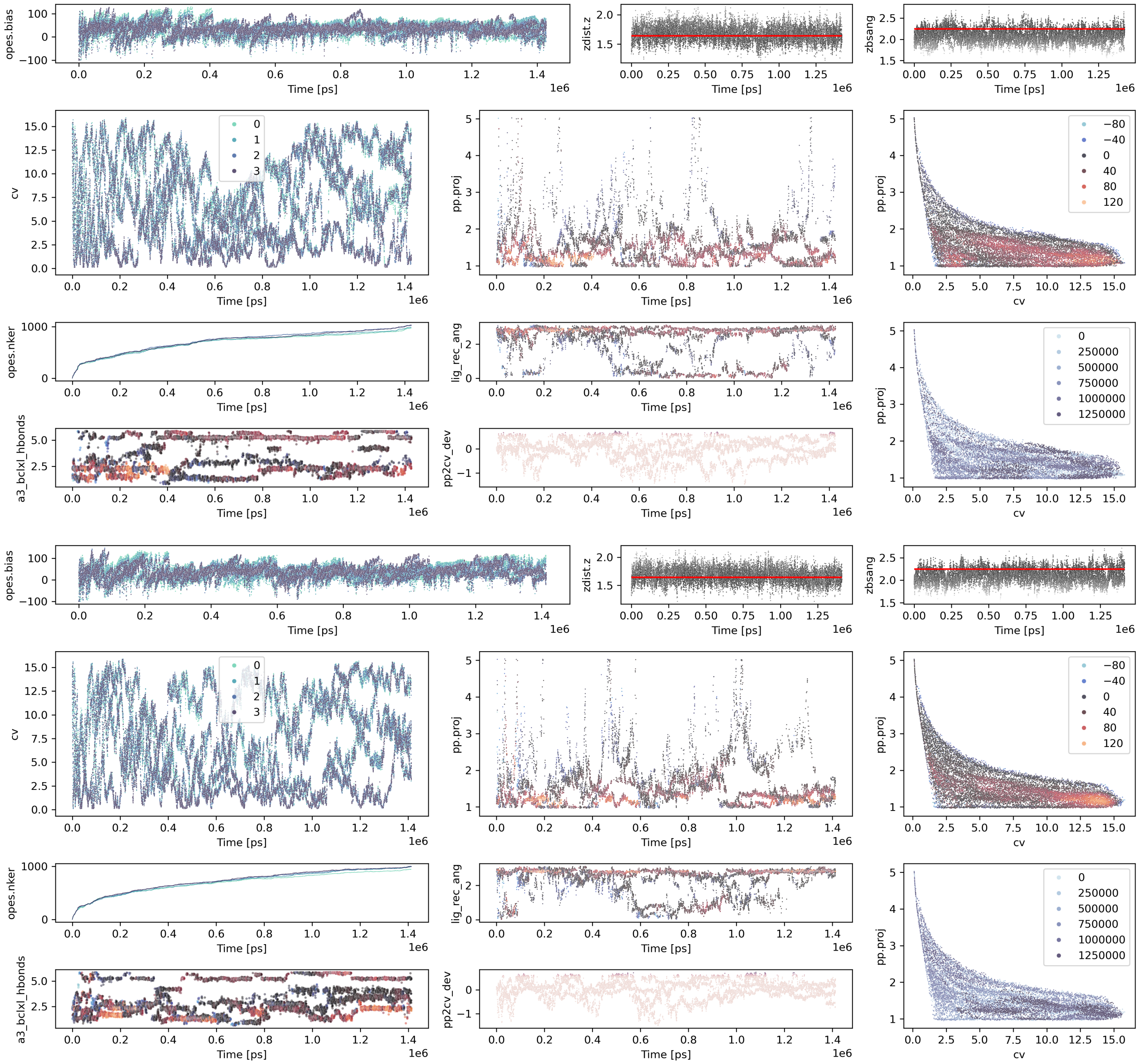


**Fig. S17. Simulation traces of four OneOPES simulations for the Bcl-xL/BidBH3 peptide system.** One set of plots per technical replicate, each set containing three rows. Top row, left to right: (1) Main opes bias applied over time colored by replicate; (2) distance of globular domain from membrane over time, red line indicates wall position; (3) Angle of the globular domain to the membrane lateral over time, red line indicates wall position. Middle row, left to right: (4) contactmap value over time colored by replicate; (5) funnel projection over time for the base replicate colored by bias; (6) Contactmap against funnel projection colored by bias in the base replica. Bottom row, left to right: (7, top) Number of kernels in the main CV opes colored by replicate; (7, bottom) Number of hydrogen bonds in the $\alpha 3$ of Bcl-xl in the base replica over time colored by the main bias; (8, top) Ligand receptor orientation over time of the base replicate colored by the main bias; (8, bottom) Deviation from the logarithmic fit between contactmap and funnel projection in the base replica over time colored by the corresponding walls bias; (9) contactmap against funnel projection colored by time in the base replica.

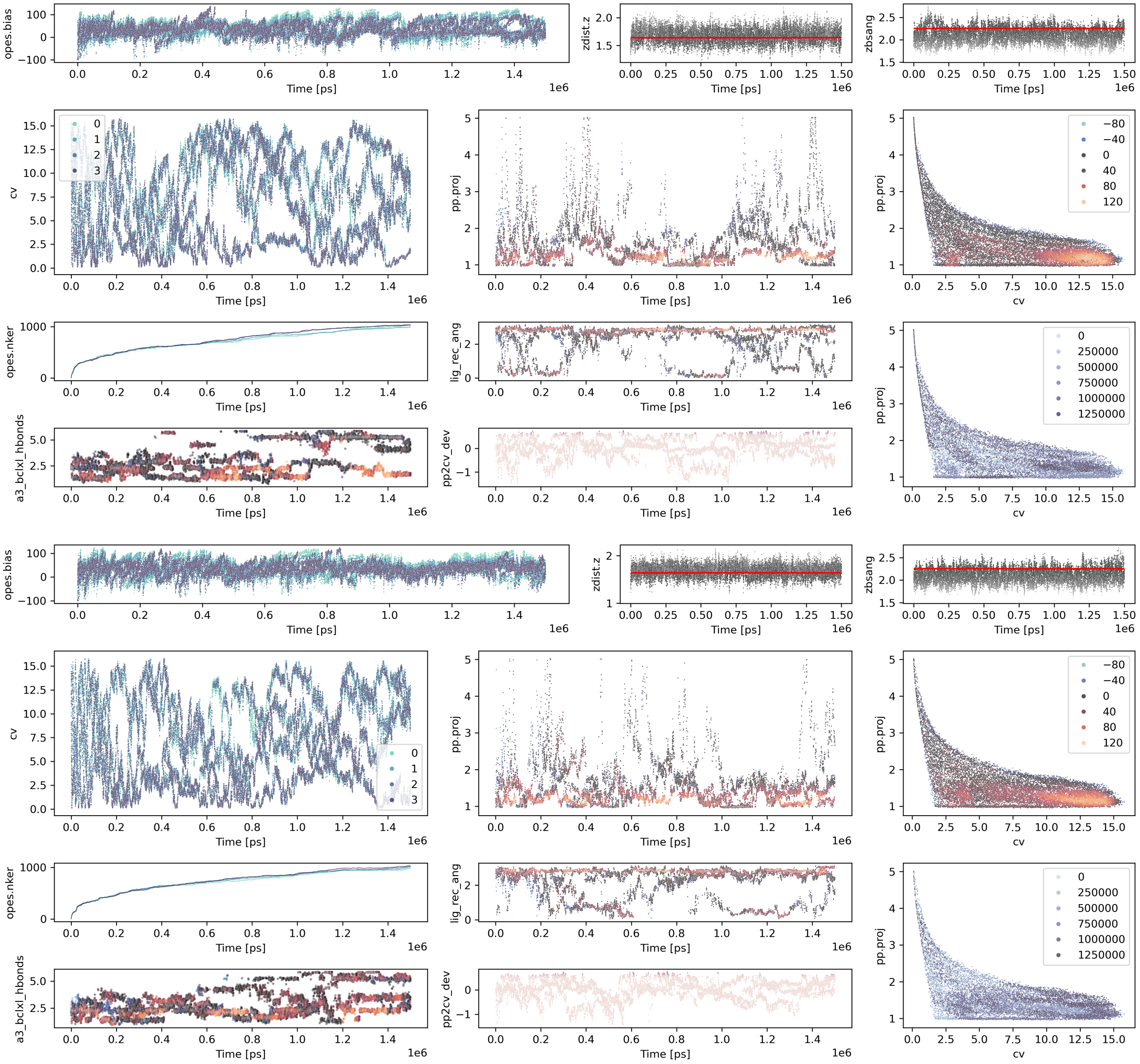

**Fig. S17. Simulation traces of four OneOPES simulations for the Bcl-xL/BidBH3 peptide system.** (continuation)
